# Comparative genomics reveals intra and inter species variation in the pathogenic fungus *Batrachochytrium dendrobatidis*

**DOI:** 10.1101/2024.01.24.576925

**Authors:** Mark N. Yacoub, Jason E. Stajich

## Abstract

The Global Panzootic Lineage (GPL) of the amphibian pathogen *Batrachochytrium dendrobatidis* (*Bd*) has been described as a main driver of amphibian extinctions on nearly every continent. Near complete genomes of three *Bd*-GPL strains have enabled studies of the pathogen but the genomic features that set *Bd-*GPL apart from other *B. dendrobatidis* lineages is not well understood due to a lack of high-quality genome assemblies and annotations from other lineages. We used Oxford Nanopore Technologies (ONT) DNA sequencing to assemble high-quality genomes of three *Bd*-BRAZIL isolates and one non-pathogen outgroup species *Polyrhizophydium stewartii* (*Ps*) strain JEL0888 and compared these to genomes of previously sequenced *Bd-*GPL strains. The *Bd*-BRAZIL assemblies range in size between 22.0 and 26.1 Mb and encode 8495-8620 protein-coding genes for each strain. A pangenome is defined as all the genes in a species including the core genome, genes found in every strain, and the accessory genome, genes found in only some strains (Brockhurst et al. 2019). To date, a comprehensive analysis identifying the core and accessory genes within *B. dendrobatidis* has not been conducted. Furthermore, while previous studies have examined the gene transcription profiles of *Bd*-GPL and *Bd*-BRAZIL strains (McDonald et al. 2020), they do not account for the genomic differences between these strains. Our pangenome analysis provides insight into shared and lineage-specific gene content and how *B. dendrobatidis* genotype affects recovery of RNAseq transcripts from different strains. We hypothesize that gene content differences exist between the *B. dendrobatidis* lineages and genomic differences, such as gene family expansions or gene sequence variation, affect alignment and enumeration of transcriptomic data when relying on a single reference genome. The pangenome analysis revealed a core genome consisting of 6278 conserved gene families, and an accessory genome with 202 *Bd*-BRAZIL and 172 *Bd*-GPL specific gene families. We discovered gene copy number differences in five pathogenicity gene families: *M36 Peptidase, Crinkler Necrosis Genes (CRN), Aspartyl Peptidase, Carbohydrate-Binding Module-18 (CBM18), and S41 Protease,* between *Bd-*BRAZIL and *Bd-*GPL strains. However, none of the five families were expanded in *Bd*-GPL compared to *Bd*-BRAZIL strains. Comparison between the *Batrachochytrium* genus and two closely related non-pathogenic saprophytic chytrids identified differences in sequence and protein domain counts. We further test these new *Bd*-BRAZIL genomes to assess their utility as reference genomes for transcriptome alignment and analysis. Our analysis examines the genomic variation between strains in *Bd*-BRAZIL and *Bd*-GPL and offers insights into the application of these genomes as reference genomes for future studies.

**Significance:** The geographically defined enzootic populations of amphibian pathogen *Batrachochytrium dendrobatidis* harbor gene context variation revealed in pan-genome analyses. Long read sequencing is required to fully capture this diversity as some recently duplicated and potential virulence gene families are undercounted in short-read only genome assemblies. This genetic variation can impact estimates of gene expression differences between strains if a single genome reference is used. It is necessary to consider the pan-genome diversity of the multiple lineages of this important amphibian pathogen and perhaps other fungal pathogens when engaging in studies of adaptation, virulence, and comparative biology of a species.

## Introduction

*Batrachochytrium dendrobatidis* (*Bd*) is a chytrid fungus and causative agent of the disease chytridiomycosis (Kilpatrick et al. 2010; Daszak et al. 1999; Longcore et al. 1999). The disease is widespread among amphibian populations and has contributed to declines on five continents (Scheele et al. 2019). The pathogen is globally distributed and genetically diverse, with strains divided into four described lineages; *Bd-*CAPE, *Bd*-ASIA-2/BRAZIL (hereafter *Bd-*BRAZIL), *Bd-*ASIA, and *Bd-*GPL (Global Panzootic Lineage) (Farrer et al. 2011; Schloegel et al. 2012; Rosenblum et al. 2013; Farrer et al. 2013; O’Hanlon et al. 2018; Byrne et al. 2019). Among these lineages, the *Bd*-GPL lineage alone is implicated in the majority of amphibian declines (Belasen et al. 2022; Becker et al. 2017; Farrer et al. 2011). The emergence of the *Bd*-GPL lineage has been suggested to be a recent expansion for the pathogen, occurring in the late 20th century (O’Hanlon et al. 2018).The other three lineages are genetically divergent from the *Bd-*GPL strains but are less widespread (Belasen et al. 2022). The genetic variation between the *B. dendrobatidis* lineages and the recent evolution of the globally proliferant *Bd*-GPL might suggest the possibility of gene expansions in *Bd*-GPL strains related to pathogenicity, allowing the *Bd*-GPL strains to outcompete strains from the enzootic lineages (Joneson et al. 2011; Farrer et al. 2017).

Despite the pathogen’s importance, little is known about gene content differences between *B. dendrobatidis* strains. The majority of genetic and genomic studies have relied on the reference genomes of two *Bd*-GPL strains, JEL423 and JAM81 (Joneson et al. 2011; Farrer et al. 2017). No contiguous genomes of *B. dendrobatidis* strains from any other lineage have been provided until now. Among the enzootic *B. dendrobatidis* lineages, *Bd-BRAZIL* has been previously compared with the *Bd-*GPL through comparative transcriptomics, regional distribution and virulence analysis (Becker et al. 2017; McDonald et al. 2020). Although strains from both lineages co-occur in South America, the *Bd-*GPL strains have been observed to be geographically unstructured and more infective towards amphibians in the Brazilian Atlantic Forest (Jenkinson et al. 2016; Greenspan et al. 2018).

Close relatives of *B. dendrobatidis* (e.g., *Homolaphlyctis polyrhiza* are saprophytic while only chytrid fungi in the genus *Batrachochytrium* are known to parasitize amphibians (Joneson et al. 2011; Martel et al. 2013; Berger et al. 1998). Expansions in peptidase, including *aspartyl*-, *M36 metallo*-, and *S41 serine-like peptidases*, have been observed in *B. dendrobatidis* compared to *H. polyrhiza* (Joneson et al. 2011; Farrer et al. 2017). A class of *Crinkler Like Necrosis* (*CRN*) genes, thought to be specific to Oomycetes, are also found in abundance in the *B. dendrobatidis* reference genome and relatives (Joneson et al. 2011; Sun et al. 2011; James et al. 2013). Chitin Binding Module 18 Domain (CBM18) containing proteins, implicated in protecting fungal pathogens against host chitinases, are similarly noted to be expanded in *B. dendrobatidis* (Abramyan & Stajich 2012; Farrer et al. 2017). Additionally, peptidases, *CRNs* and *CBMs* appear to be up-regulated during *B. dendrobatidis* infection compared to when the fungus is grown on media, further implicating their role in pathogenicity (Rosenblum et al. 2012; Ellison et al. 2017). Recently a newly described species, *Polyrhizophydium stewartii* has been identified and classified as a closer saprophytic relative of *B. dendrobatidis* than *H. polyrhiza* (Simmons et al. 2021; Amses et al. 2022). The discovery of *P. stewartii* provides a resource to improve understanding of the transition towards pathogenicity in chytrids. Genomic comparisons between *P. stewartii* and *B. dendrobatidis* will expand upon previous comparisons between *B. dendrobatidis* and saprophytic chytrids.

Pangenomes are valuable tools to elucidate the genomic variation within a population by classifying genes that are present in every strain (core gene families) or present in only certain lineages of the species (variable gene families) (Brockhurst et al. 2019). The mechanisms that allow *Bd*-GPL to globally proliferate while strains from other lineages remain endemic is not understood. Competition between *B. dendrobatidis* lineages, differences in virulence between *Bd*-GPL and endemic lineages, and unique genomic recombination in *Bd*-GPL have been demonstrated (Belasen et al. 2022; Jenkinson et al. 2016; Farrer et al. 2011). However, no analysis has been conducted to study the pangenome of *B. dendrobatidis* particularly concerning the conservation of pathogenicity genes; *aspartyl*-, *M36 metallo*-, and S*41 serine-like peptidases, Crinkler Like Necrosis genes* (*CRN*), and Chitin Binding Module 18 Domain (*CBM18*) containing proteins. We hypothesized that the *Bd*-GPL lineage will possess unique copies or orthologs of pathogenicity genes (*M36, S41, CRN, ASP,* or *CBM18*) that are absent within strains from the enzootic *Bd*-BRAZIL lineage. Furthermore, we suspect that genomic differences, such as copy number differences of genes or sequence level variation of orthologous genes, may inflate the transcriptomic differences between strains, a factor not accounted for in previous gene expression analysis (McDonald et al. 2020). To test these hypothesis we utilized Oxford Nanopore Technologies (ONT) sequencing of three *Bd-*BRAZIL strains and the *P. stewartii* strain *Ps* JEL0888 to aid in the establishment of a *B. dendrobatidis* pangenome. We then utilize the new *B. dendrobatidis* reference genomes to test how RNAseq transcript abundance counts differ compared to when they are aligned to the previous reference genome (GCA_000149865). Our assembly of *Bd*-BRAZIL strain CLFT044 suggests an improvement on previous assemblies, possessing three telomere-to-telomere scaffolds, 15 scaffolds with a telomeres on only the 3’ end and 11 scaffolds with telomere’s on only the 5’ end. Furthermore, the BUSCO genome completeness estimates indicate that all three *Bd*-BRAZIL genomes; CLFT044, CLFT067 and CLFT071, are of similar quality to the previous *Bd*-GPL assemblies; JEL423, JAM81 and RTP6 (ranging from 88.2% to 88.9% completeness), suggesting their utility in future genomic analysis. We present these new genomes to elucidate the genome content variability between *Bd-*GPL and *Bd-BRAZIL* strains.

## Results

### Assembly quality of the Bd-BRAZIL genomes are comparable to that of the Bd-GPL reference genomes; JEL423, JAM81 and RTP6

Our assembled *Bd-*BRAZIL genomes were assessed to be of similar quality to the three published GPL genomes in measures of total contig count, N50, and BUSCO completeness for genomes and gene annotations (**Supplementary table 1**). After assembly the *Bd-*BRAZIL genome assemblies composed 79, 86, and 85 contigs for CLFT044, CLFT067, and CLFT071 respectively with total lengths between 22Mb and 26Mb. BUSCO assessment of the assemblies concluded similar scores with average completeness for all *Bd-*BRAZIL genomes at 89.9% against the fungi_odb10 gene sets, slightly higher than the reference genome JEL423 at 88.2% completeness. Similarly, the BUSCO completeness scores for the *Bd*-BRAZIL genome annotations ranged from 91.7% to 92% and were higher on average than the *Bd*-GPL genomes JEL423, JAM81, and RTP6 (81.2%, 88.8%, and 92.1% respectively). The assembly of *Ps* JEL0888 contained 291 scaffolds with an average BUSCO completeness of 81.4% for the genome assembly and 87.9% for the annotation. The *Ps* JEL0888 genome assembly size was 31Mb, significantly larger than the *B. dendrobatidis* assemblies which ranged from 22-26Mb. The *H. polyrhiza* assembly (GCA_000235945.1) was far less contiguous and complete than those of the other chytrids, being assembled exclusively from short-read sequencing data. The *H. polyrhiza* assembly possesses 11986 scaffolds and a BUSCO completeness of 80.9. Furthermore, its largest scaffold was only 227.1kb long, far shorter than the largest scaffolds of the other chytrid assemblies.

### *B. dendrobatidis* is expanded in genome size and TE content compared to *H. polyrhiza but* reduced compared to *P stewartii*

*Bd-*BRAZIL and *Bd-GPL* strains were overall similar in genic space and transposable element (TE) content (**Figure 1**). However, the *Bd*-BRAZIL strain, CLFT044 possesses a greater abundance of Unclassified elements (13958 copies/6.12Mbp), Long Terminal Repeat (LTR) elements (977 copies/1Mbp) and DNA transposons (1409 copies/1.12Mbp) compared to the other *B. dendrobatidis* strains, which contain an average content of 12958 copies/4.23Mbp, 686 copies/562Kbp and 1250 copies/800Kbp for Unclassified, LTR, and DNA elements respectively. Overall the TE content of *B. dendrobatidis* was expanded compared to that of *H. polyrhiza* which possessed only 147 copies/19Kbp LTR elements and 271 copies/38Kbp DNA transposons. Our results support previous findings that *H. polyrhiza* has a reduced genome size and TE content compared to *B. dendrobatidis* (Farrer et al. 2017).

**Figure 1:**
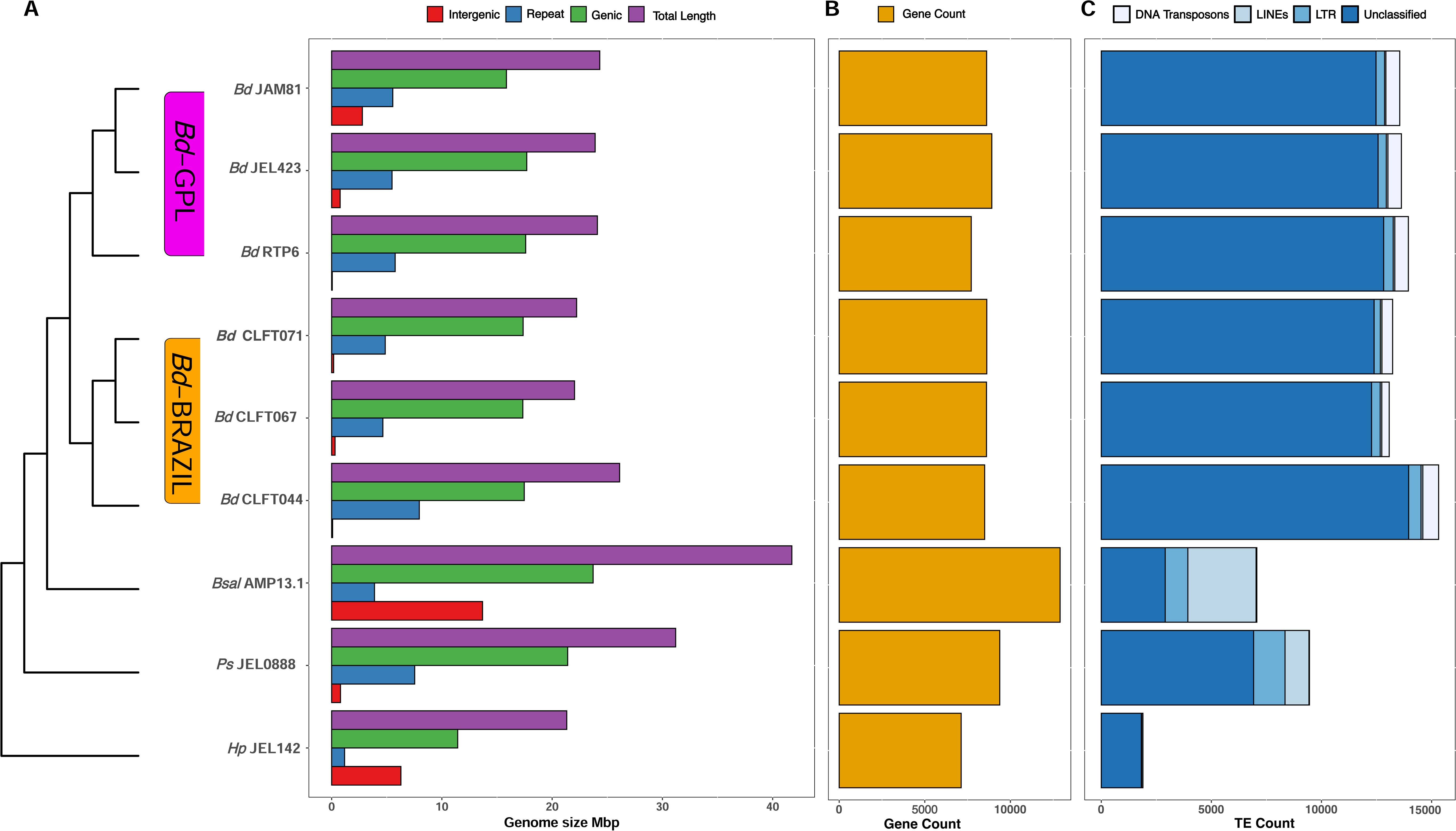
Variation in the overall total length, gene counts, repeat TE counts between *B. dendrobatidis* strains, *B. salamandrivorans*, *H. polyrhiza*, and *P. stewartii*. (Left) A barplot is shown depicting the length in Mbp of overall genome size (purple), genic space (green), non-repeat intergenic space (red), and TE content (blue). (Middle) Gene counts in the genomes are shown in the gold barplot. Scale is in the number of genes. (Right) TE counts are represented by the blue barplot. TEs were classified as either DNA, LINE, or LTR elements.

The gene counts found in all *B. dendrobatidis* strains were higher than that of *H. polyrhiza,* however *P. stewartii* possesses a greater number of genes than any of the *B. dendrobatidis* strains *B. dendrobatidis* gene counts ranged from 7712 to 8914 genes, *P. stewartii* possesses 9378 genes and *H. polyrhiza* possesses 7123 genes. All other species were dwarfed in gene count compared to *Batrachochytrium salamandrivorans*, a close relative of *B. dendrobatidis* and pathogen of salamanders, which contains 12900 genes. TE sequences can be found in the github repository for this manuscript: **DOI 10.5281/zenodo.15070156.**

TE expansion has been suggested as a driving force in the acquisition of pathogenicity genes in *Batrachochytrium* (Wacker et al. 2023). We identified expansions of LTR, Long interspersed nuclear element (LINE), and DNA TEs in *B. dendrobatidis* and *B. salamandrivorans* compared to *H. polyrhiza*’s TE content*. P. stewartii* was expanded in LTRs (1832 copies/2.04Mbp) compared to *B. dendrobatidis*, however its genome contains fewer DNA TEs (132 copies/68Kbp) than any of the *B. dendrobatidis* strains. All *B. dendrobatidis* strains are expanded in DNA TEs compared to the other species including *B. salamandrivorans* (430 copies/84.2Kbp). Although a Pearson’s correlation test revealed that TE content (bp) was not significantly correlated with genome size between species (R=0.11, p=0.77), TE content was significantly correlated with genome size differences between *B. dendrobatidis* strains (R=0.94, p=0.0059).

### Synteny is conserved along the 10 largest scaffolds between CLFT044 and the GPL reference genomes

Conserved syntenic regions between the longest 10 scaffolds in *Bd*-BRAZIL strain CLFT044 were compared with their homologs in *Bd*-GPL strains JEL423 and RTP6 (**Figure 2**). The 10 largest scaffolds in CLFT044 were present in the RTP6 and JEL423 assemblies. Scaffold_5 and Scaffold_9 appear to be split in CLFT044, being combined as DS022301.1 and QUAD01000002.1 in JEL423 and RTP6 respectively. Scaffold_5 contains one telomere on the 5’ end while Scaffold_9 does not possess telomeric sequences. This suggests completeness on the 5’ end of this region while the center and 3’ ends of the chromosome were not contiguously assembled.

**Figure 2:**
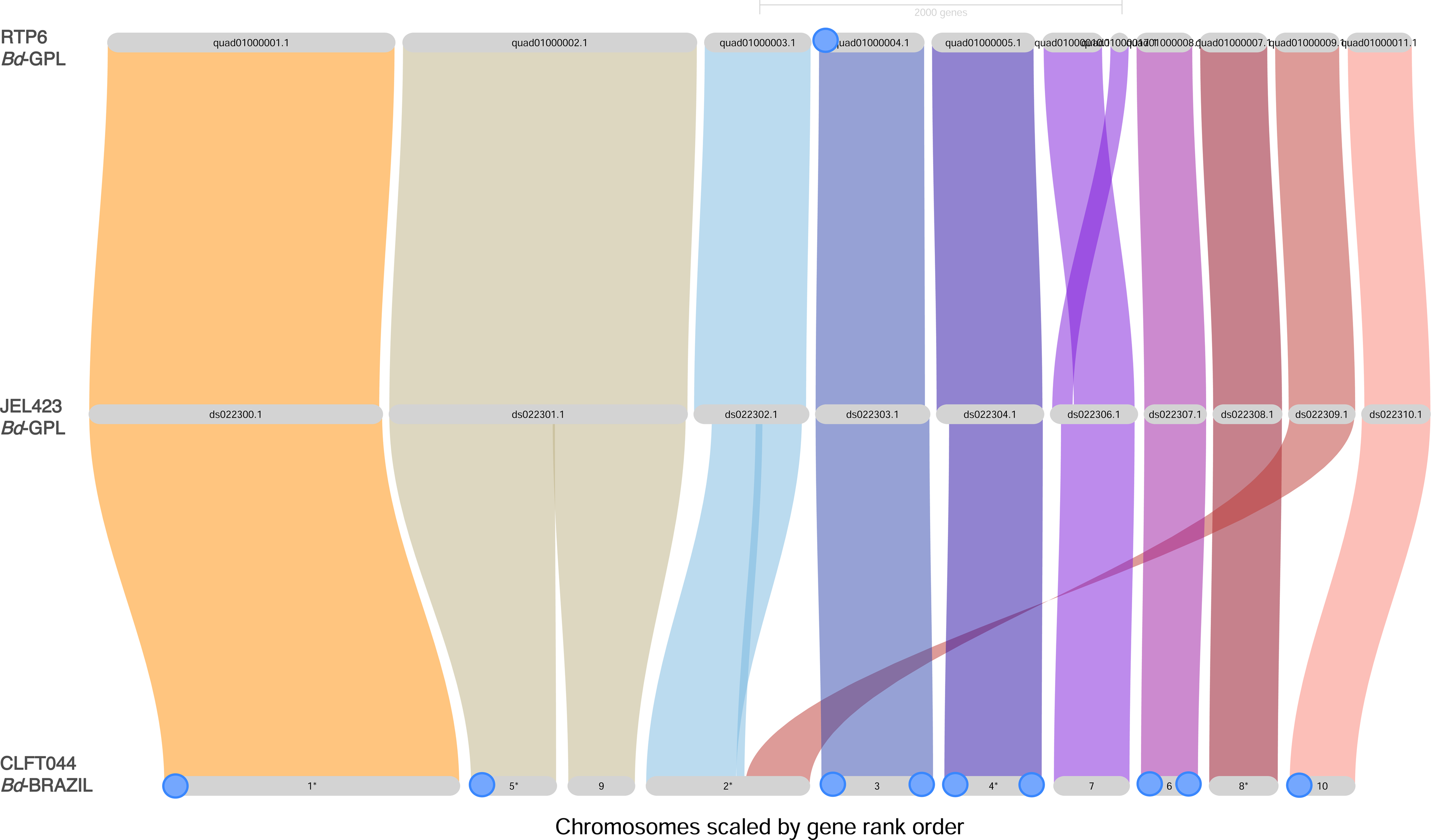
A Riparian plot generated by GENESPACE v1.2.0 depicting synteny between scaffolds in CLFT044, JEL423, and RTP6. Orthology was used to visualize and compare synteny between the 10 longest scaffolds in CLFT044 and their homologs in RTP6 and JEL423. Genomes are organized by row and scaffolds are organized by column. Linkages are color coded by their respective scaffold in the reference genome JEL423. Telomeres are depicted as blue circles on the scaffold labels.

Scaffold_2 in CLFT044 represents a more contiguous assembly, being a merge of the two contigs DS022302.1 and DS022309.1 in JEL423. ONT reads from CLFT044 spanned the entire Scaffold_2 region, indicating the validity of this merged scaffold compared to other assemblies.

Scaffold_3, Scaffold_4, and Scaffold_6 in CLFT044 all contained forward and reverse telomeres repeats based on the telomere pattern search results suggesting their status as complete chromosome assemblies. Although Scaffold_4 in CLFT044 is likely a complete chromosomal assembly, its homolog extends farther on the 5’ end in JEL423 and RTP6. This difference in length could be due to genome size differences between strains rather than assembly quality.

### *Bd-*BRAZIL and *Bd-*GPL possess many lineage specific gene families

Analysis of shared orthogroups between *B. dendrobatidis* and closely related species (**Figure 3A**) reveals 348 gene families that are distinct to the *Batrachochytrium* genus (all *B. dendrobatidis* strains and *B. salamandrivorans* but absent in the saprophytic chytrids. Additionally, 435 gene families were unique to *B. dendrobatidis*, being present in all *B. dendrobatidis* strains but absent in the other chytrid species. Gene Ontology (GO) enrichment analysis on the JEL423 genes found in the pathogen specific gene families indicates pathogen specific expansions in *Metallopeptidase* gene families (**Supplemental figure S2**). Our Computational Analysis of Gene Family Evolution (cafe), which identifies expansions and contractions of gene families between genomes, revealed significant expansions of orthogroups between *H. polyrhiza*, *P stewartii*, and *B. dendrobatidis* (**Supplemental Figure S3**). We observed 148 expanded and 147 contracted gene families in all *B. dendrobatidis* strains compared to *P. stewartii. Bd*-BRAZIL exhibited an additional 153 expanded and 37 contracted gene families unique to this lineage while 63 unique expansions and 22 contractions were observed in *Bd*-GPL.

**Figure 3:**
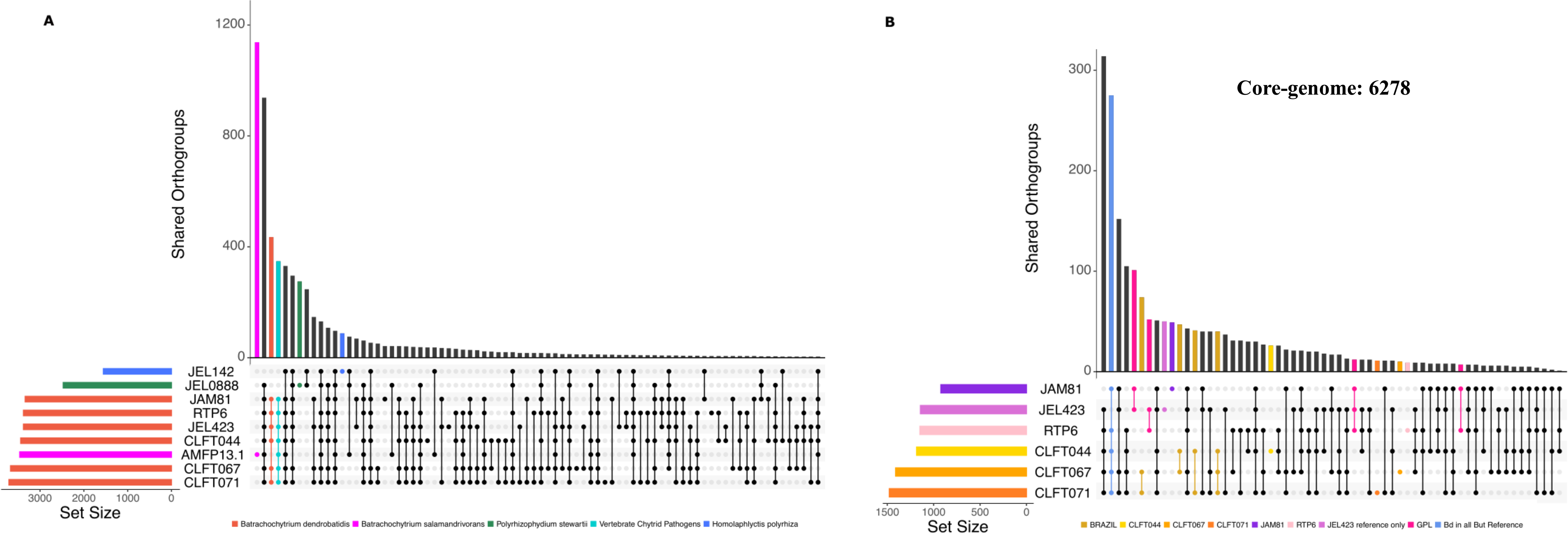
Orthofinder Pangenome analysis between chytrid species and between *B. dendrobatidis* strains visualized as Upset plots. (A) Upset plot depicting Orthofinder results for six *B. dendrobatidis* strains and related species *B. salamandrivorans, P. stewartii, and H. polyrhiza* and (B) between *B. dendrobatidis* strains only. Vertical bars display the counts of gene families in each group while the dots indicate the strains present in each group. Only accessory and singleton gene families are shown. Gene families found in a single strain are indicated as a single dot and colored by strain.

Orthofinder results between *B. dendrobatidis* strains indicate that there are 6278 core gene families in the *B. dendrobatidis* pangenome (**Figure 3B**). We identified 1,934 accessory gene families that were present in two or more strains and 160 singletons present in single strains. Among the accessory gene families, we find 202 to be distinct to the *Bd*-BRAZIL strains while 172 are unique to *Bd*-GPL. The singletons are unequally divided among strains with strains JEL423 and JAM81 having the highest number of unique gene families (51 and 50 respectively) while the remaining strains each possessed only 10-30 singleton gene families. We identified a gene family of putative Meiotically up-regulated gene 113 (mug113) proteins found in other fungi and in the *Bd-*BRAZIL lineage but absent in *Bd*-GPL. Among the *Bd*-GPL specific gene families were many proteins of unknown function that were absent in all *Bd*-BRAZIL strains. Interestingly the reference genome strains JEL423 and JAM81 lacked 105 gene families that are present in all the other *B. dendrobatidis* genomes (**Supplementary figure S4**).

### Pathogenicity genes vary in count and sequence between *B. dendrobatidis* strains and other chytrids

Our HMMsearch analysis of the five pathogenicity genes revealed differences in copy number and sequence diversity of these gene families among *B. dendrobatidis* strains and sister species (**Figure 4**). Among the putative pathogenicity genes, all *B. dendrobatidis* strains were expanded in copy number for *S41 Peptidase*, *ASP protease*, *CBM18*, and *CRN* relative to *B. dendrobatidis*’s saprophytic relatives, *H. polyrhiza* and *P. stewartii*. *CRN* had the highest observed expansion in *B. dendrobatidis* compared to its saprophytic relatives which ranged from one to six copies in *H. polyrhiza* and *P. stewartii* and 27-108 copies in the *B. dendrobatidis* strains. Although we observed differences in pathogenicity gene content between *B. dendrobatidis* strains, the gene families were not consistently expanded in *Bd*-GPL over *Bd*-BRAZIL. We compared the pathogenicity gene counts detected in long-read genomes of CLFT044, CLFT067, CLFT071 and JEL423 with the counts found in the annotated genomes assembled from only Illumina short read genomes for the same strains to determine whether sequencing technology plays a significant role in capturing Pathogenicity gene diversity (**Supplementary figure S5**). While counts for the *SWEET* and *Adenylate kinase* gene families were consistent between Illumina and Long-read genomes, the pathogenicity genes were consistently under-represented in the Illumina assemblies compared to the long-read genomes for the same strain. We confirmed that short, Illumina reads from the *B. dendrobatidis* strains indiscriminately align to highly similar and multi-copy homologs in their ONT assemblies. This suggests that de novo assembled genome from Illumina-only data collapse expanded gene families into fewer representatives. We previously hypothesized that the *Bd*-GPL lineage will possess unique expansions of pathogenicity genes such as *M36* or *CBM18*, that are absent within strains from the enzootic *Bd*-BRAZIL lineage. To test this hypothesis we examined the presence of all orthologs in these gene families across the *B. dendrobatidis* pangenome to determine if any specific copies were lineage specific. Using cblaster we resolved *M36* and *CBM18* genes into orthologous loci using flanking syntenic genes. To classify M36 and CBM18 genes as core, variable, or singleton in the *B. dendrobatidis* pangenome, we grouped homologous sequences from multiple strains based on their co-location, representing each group by a single ortholog from one strain. We searched with cblaster to find these representative orthologs in the *B. dendrobatidis* pangenome and determine the distribution of core and variable *CBM18s* and M36*s* in the *B. dendrobatidis* pangenome. If the core or variable *M36/CBM18* was found in JEL423, the representative was named with the JEL423 copy, otherwise it was named for the representative from another *B. dendrobatidis* strain. Prior to collapsing duplicate orthologs there were a total of 210 *M36* and 91 *CBM18* copies in all six *B. dendrobatidis* strains. Collapsing the orthologous copies from separate strains resulted in 59 and 20 unique orthologs of *M36* and *CBM18* respectively.

**Figure 4:**
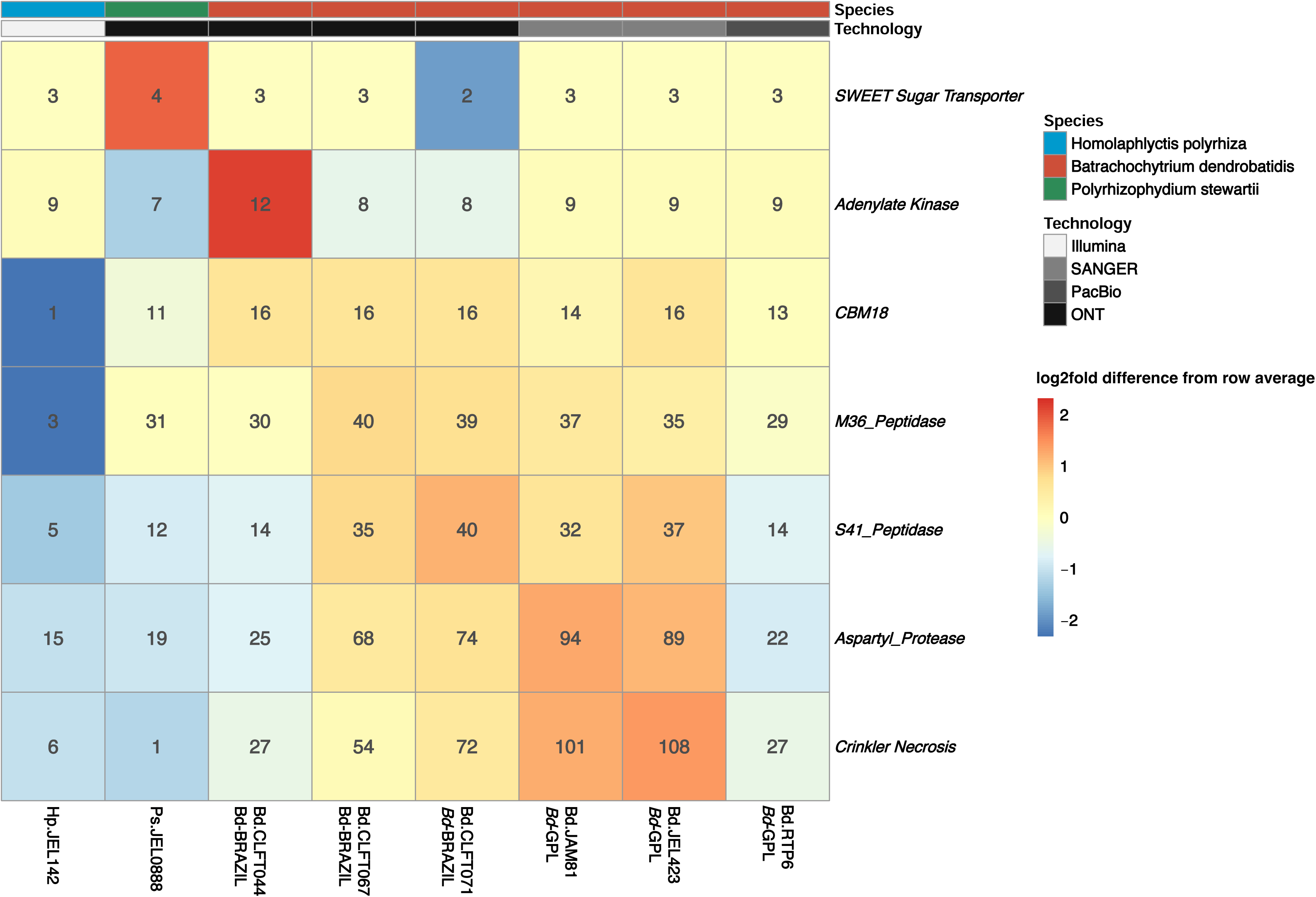
Copy number variation for pathogenicity genes *CBM18, M36, S41, ASP*, and *CRN* between *B. dendrobatidis* and saprophytic chytrids. A heatmap is shown depicting the copy number variation of pathogenicity genes (SWEET and Adenylate Kinase included as controls). The colors of the tiles represent log2fold difference from the average and are normalized per gene family with red and blue indicating high and low counts respectively. The numbers within the tiles indicate the number of homologs of each gene family in each genome. The sequencing technology used to produce each genome is shown by the gray-scale bar while the top color bar indicates the species for each strain. *B. dendrobatidis* lineages (*Bd*-BRAZIL or *Bd*-GPL) are indicated beneath their respective strain names.

Phylogenetic analyses of the *M36* orthologs revealed ancestral and *B. dendrobatidis* exclusive clades of this gene family (**Figure 5A**). Despite similar counts of *M36* genes between *P. stewartii* and *B. dendrobatidis*, we observed sequence level variation that segregated the *B. dendrobatidis M36* orthologs from the *P. stewartii* orthologs. We defined the ancestral M36 clade as the clade of *M36* orthologs that contained members of *M36* found in all genomes and species while the *B. dendrobatidis* specific clade contains orthologs found only in *B. dendrobatidis*. Furthermore, we identified a previously unknown clade of *M36* genes unique to *P. stewartii. B. salamandrivorans* ’s *M36* repertoire is expanded compared to *B. dendrobatidis,* with two clades of *B. salamandrivorans* specific *M36* clades sister to the *B. dendrobatidis* specific clade (**Supplementary figure S6**). *B. salamandrivorans* and *B. dendrobatidis* possess similar counts of *M36* genes in the ancestral *M36* clade with three copies in *B. salamandrivorans* and six in *B. dendrobatidis*.

**Figure 5:**
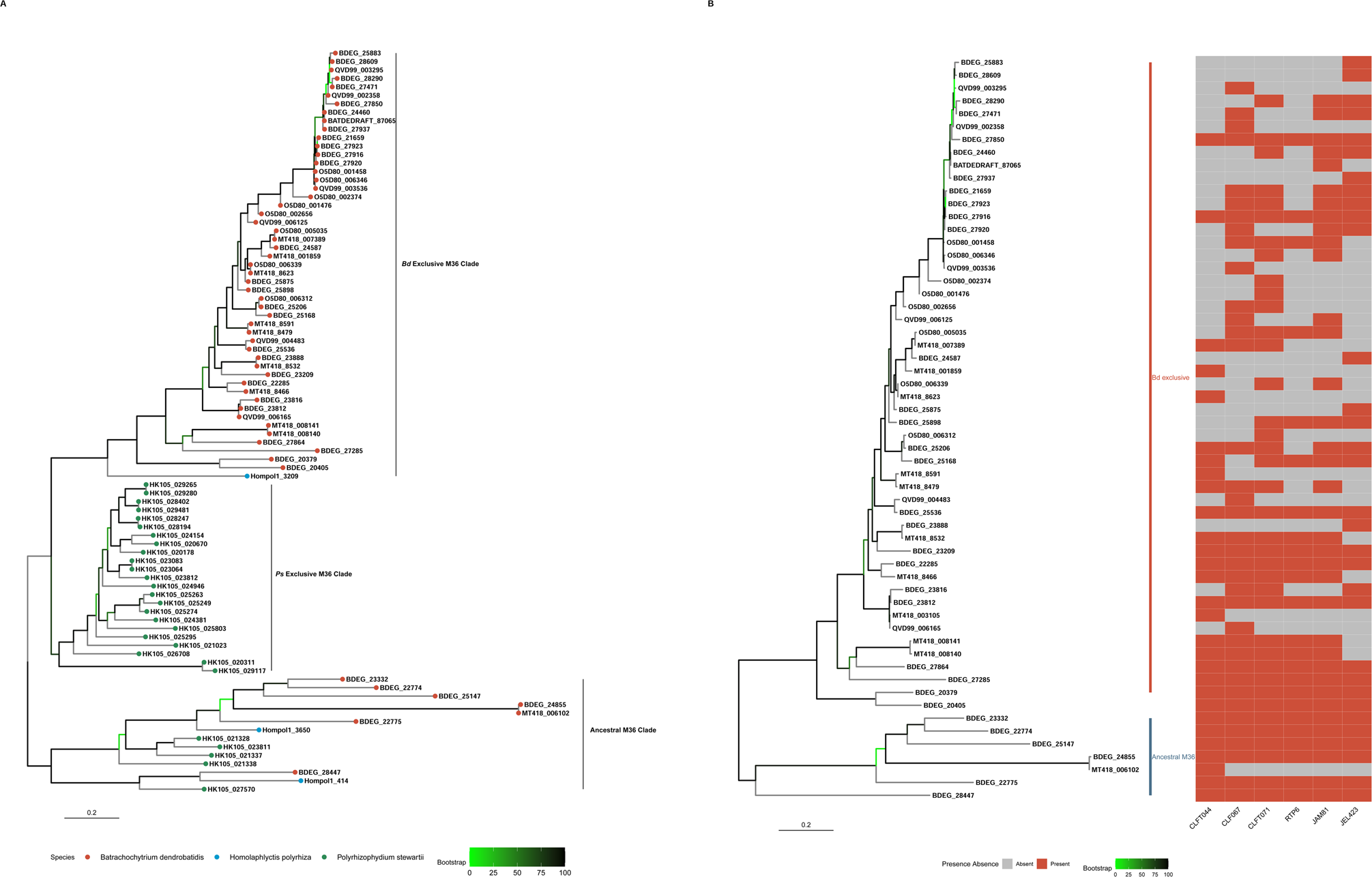
Phylogenetic relationships and distribution of *B. dendrobatidis* exclusive *M36* loci and those from saprophytic chytrids. (A) Maximum likelihood phylogeny inferred by IQTREE of the unique *M36* genes in *B. dendrobatidis*, *H. polyrhiza,* and *P. stewartii* is depicted here. Duplicate orthologs are collapsed by results from cblaster. Tips are color coded based on the species that M36 homolog is found in. The clade of *M36* genes exclusive to *B. dendrobatidis* is labeled as “*Bd* exclusive” and the ancestral clade containing representative *M36* genes from all species is labeled as “Ancestral”. Bootstrap support values are color-coded at the nodes of the phylogeny with green nodes indicating branches with low support. (B) Phylogeny of unique M36 orthologs and their presence/absence in the six *B. dendrobatidis* genomes. A reduced *M36* tree from figure 5A, focused on *M36* from only *B. dendrobatidis.* The heatmap represents the percentage of Illumina genomes in each lineage that contain that M36 homolog. The ancestral M36 clade is labeled. A scale bar is included which indicates nucleotide substitutions per site.

We searched with cblaster for the newly identified *M36* ortholog clusters against the six Long-read *B. dendrobatidis* genomes to establish the conservation of these *M36* orthologs across the *B. dendrobatidis* strains (**Figure 5B**). We identified 15 core *M36* orthologs conserved in all six genomes. While we did not identify any *Bd*-GPL specific *M36* orthologs we discovered two *Bd*-BRAZIL specific *M36* clusters that were present in two-three *Bd*-BRAZIL strains but absent in all *Bd*-GPL. Additionally, we reveal the presence of nine *M36* singleton gene copies that are present in only a single strain. We analyzed the region of MT418_006102, a singleton *M36* from *Bd-*BRAZIL strain CLFT044, and compared its syntenic structure to its closest relative BDEG_24855, a core *M36* from the ancestral clade. We discovered that the flanking region of MT418_006102 was duplicated and downstream of the BDEG_24855 syntenic cluster in CLFT044 (**Supplementary figure S7**).

Annotation error has likely contributed to the *M36* count variation between *B. dendrobatidis* strains. The analysis with cblaster revealed the JEL423 gene BDEG_27858 to be orthologous with a pair of tandem *M36* genes in all other strains (**Supplementary figure S8**). The combined length of the *M36* pair in other strains was equal to the length of BDEG_27858 further suggesting the genes were erroneously merged during annotation of the reference genome. One of the *M36* loci in this region was likewise fragmented in JAM81 resulting in three *M36* genes rather than two.

We identified variation in the domain content of *CBM18* genes between *B. dendrobatidis*, *P. stewartii*, and *H. polyrhiza* (**Figure 6A**). *B. dendrobatidis* and *H. polyrhiza* both possess only one copy of Tyrosinase containing CBM18 proteins while *P. stewartii* contains three that are co-localized, possibly a result of tandem duplication. Lectin-like CBM18 proteins were expanded in *B. dendrobatidis* compared to either of its relative species, with only two present in *P. stewartii*, none in *H. polyrhiza,* and 10-14 orthologs in each *B. dendrobatidis* strain. Additionally, we discovered differences in CBM18 Domain counts within homologs between the *B. dendrobatidis* strains (**Figure 6B**). We found that most *CBM18* homologs (except for BDEG_23733) varied in CBM18 domain count between different strains. One example displaying such diversity was BDEG_21734 which contains five domain copies in all *Bd*-GPL strains and four per *Bd*-BRAZIL. Furthermore, homologs of BDEG_20255 which, although present and syntenic in all *B. dendrobatidis* genomes, varied significantly in CBM18 domain counts with three to five domains per strain (**Supplementary figure S9)**.

**Figure 6:**
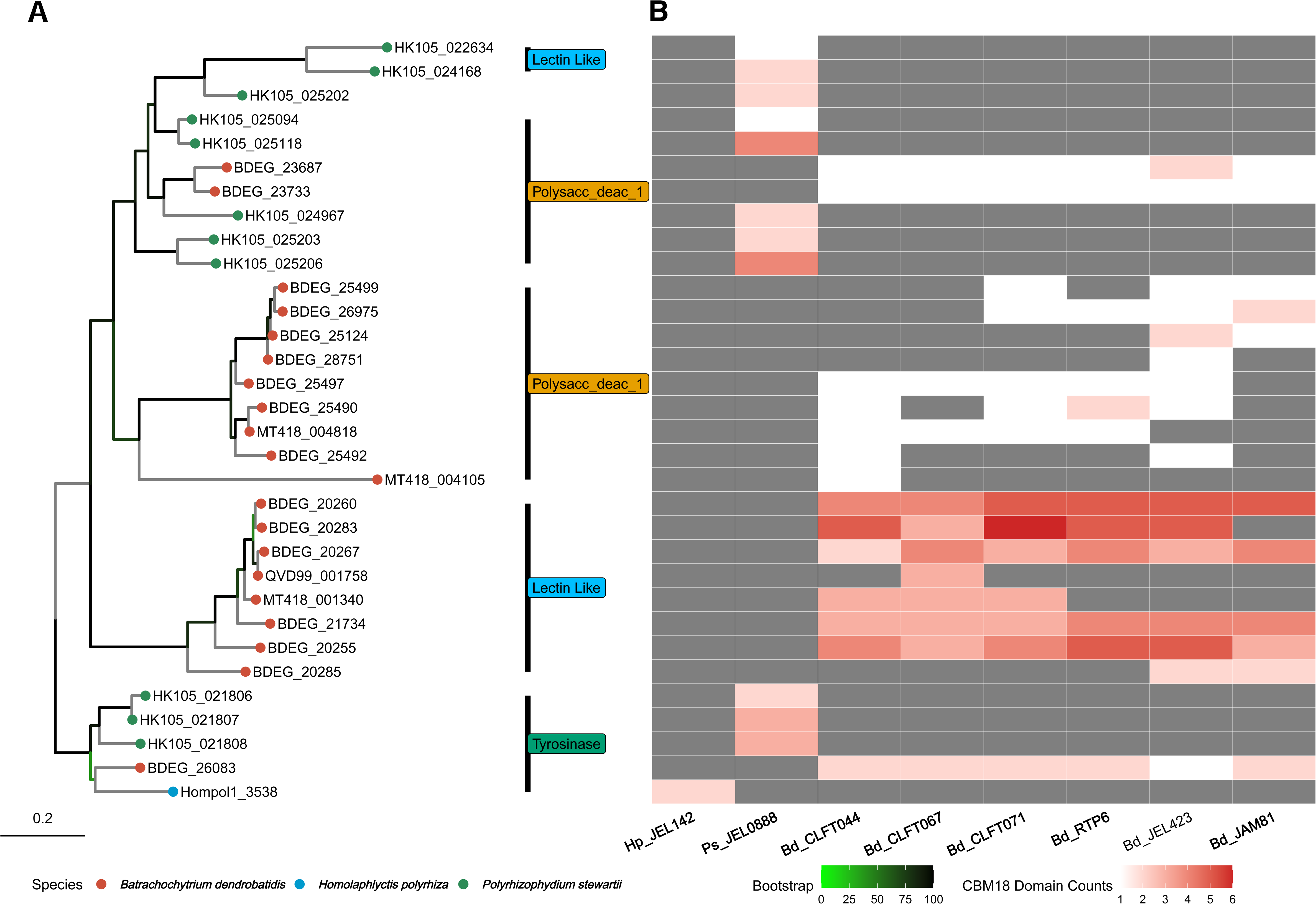
Homologous *CBM18* genes display variable domain counts between *B. dendrobatidis* strains. Maximum-likelihood *CBM18* whole gene phylogeny inferred by IQTREE with duplicate orthologs collapsed based on blaster results from *B. dendrobatidis, P. stewartii*, and *H. polyrhiza.* Clade labels define genes within each clade by class; Tyrosinase, Deacetylase, and Lectin-like. Tree tips are coded by species. In association with the tree tips, a heatmap depicts domain counts for that CBM18 homolog. Columns of the heatmap indicate the genomes of *H. polyrhiza, P. stewartii,* and the long-read *B. dendrobatidis* genomes. White colors indicate low-counts of CBM18 domains for the homolog in that genome. Red indicates that strain has a high count of CBM18 domains in that protein. Gray indicates the absence of the entire CBM18 homolog. Bootstrap support values are color-coded at the nodes of the phylogeny with green nodes indicating branches with low support. A scale bar is included to indicate nucleotide substitutions per site.

### Aligning transcripts from *Bd-*BRAZIL strains to JEL423 may under or over-estimate gene expression for some genes

Our RNA-seq analysis with HISAT2 revealed that transcripts from Bd-BRAZIL strains aligned to the CLFT044 genome at an average mapping percentage of 97.52% and to the JEL423 genome at 96.85%. We found that overall, the Transcripts per kilobase million (TPM) ratios for CLFT044 transcripts when aligned to CLFT044 vs. JEL423 varied little, with the average TPM ratio at approximately 1 (0.993) (**Figure 7B**). Despite the largely consistent average mapping percentage, we found that the CLFT044 transcripts from 145 single copy genes are under-represented (TPM ratio > 1.2) and 279 are over-represented (TPM ratio < 0.79) when aligning to the JEL423 genome (**Figure 7C**). Gene length variance, percent ID, and number of secondary blast hits between JEL423 and CLFT044 were significantly correlated to TPM variance, however we were unable to determine a conclusive cause for this variance (**Supplementary figure S10**).

**Figure 7:**
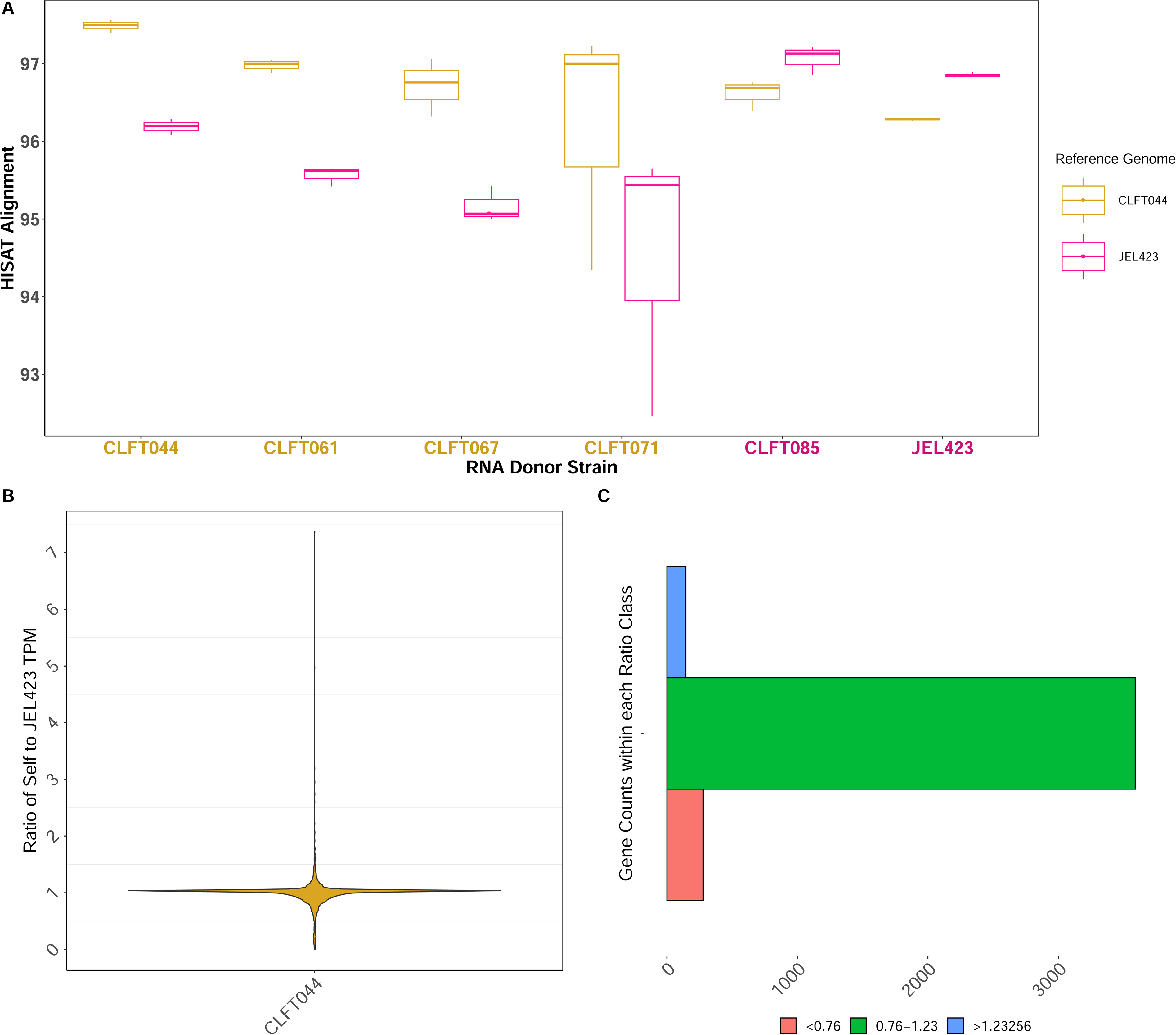
*B. dendrobatidis* reference genome affects recovery and alignment of RNA-seq transcripts. Alignment of *Bd*-BRAZIL strain transcripts to CLFT044 and JEL423. (A) The percentage of aligned transcripts from four *Bd*-BRAZIL strains and two *Bd*-GPL strains to the reference genomes CLFT044 and JEL423 is shown. X-axis indicates the RNA-seq donor strain. Gold x-axis labels indicate the RNA-seq donor strains from the *Bd*-BRAZIL lineage and purple labels are strains from *Bd*-GPL. Boxplots are color coded by reference genome. (B) A violin plot is shown depicting the TPM ratio for CLFT044 RNA-seq data when aligned to the CLFT044 genome versus aligning to the JEL423 genome for the single-copy orthologous genes. Each gold point represents a single copy ortholog gene and the y-axis represents the ratio of TPM when aligned to CLFT044 over JEL423. (C) The number of genes with TPM ratios below (red), within (green), and above (blue) one standard deviatoin of “one”. Transcript counts of genes in the red group are underestimated when aligned to the previous reference genome while genes in the blue group are over-estimated. Tanscripts from genes within the green class are accurately counted regardless of reference genome.

Differential expression analysis using RNA-seq data from a previous study (McDonald et al. 2020) revealed 1083 DEGs between the *Bd*-BRAZIL and *Bd*-GPL lineages using JEL423 as the reference genome. While Gene Ontology (GO) terms were only available for 538 of these DEGs, genes up-regulated in *Bd*-GPL compared to *Bd*-BRAZIL included nine genes with metallo-peptidase activity (including nine homologs of *M36* identified and in this study) and nine genes with Serine peptidase activity (S41). 18 DEGs up-regulated in *Bd*-BRAZIL compared to *BD*-GPL included GO terms for aspartic peptidase activity (ASP). Among the total DEGs, 89 genes were specific to the *Bd*-GPL lineage, 43 were among the single-copy orthologous genes identified as over or under-represented when aligning RNA to the JEL423 reference genome (**Supplementary figure 11A**). Aligning the same RNA-seq data to the CLFT044 genome resulted in 788 DEGs. 31 of the Differentially Expressed Genes (DEGs) were *Bd*-BRAZIL specific genes and 49 were genes with reference dependent transcript counts (**Supplementary figure 11B**).

## Discussion

*B. dendrobatidis* is the causative agent of chytridiomycosis in amphibian populations around the world, including North and South America (Longcore et al. 1999). However, the pathogen is not a monolith, as strains from different lineages exhibit variable distribution and virulence on their amphibian hosts (Dang et al. 2017; Rosenblum et al. 2013). Little is known about the functional differences that drive pathogenicity variance between the *Bd-*GPL and *Bd-*BRAZIL lineages; therefore, these new *Bd*-BRAZIL genomes represent a valuable dataset through which to examine this variance. Although genome expansions and TE invasions have been implied as a driving force in hypervirulence between some strains of plant pathogenic fungi (Grandaubert et al. 2014) TE content was not correlated with genome size between chytrid species. In contrast, content of Gypsy, Copia, DNA transposons and unclassified elements were all highly associated with genome size differences between *B. dendrobatidis* strains (R=0.94, p=0.0059). This suggests that the variation in genome size within the *B. dendrobatidis* species may be at least partly explained by TE expansions. Despite this correlation, we did not detect an abundance of TEs in *Bd*-GPL strains compared to *Bd*-BRAZIL even when accounting for unclassified TE families. Similarly, our counts of pathogenicity genes between *B. dendrobatidis* strains reveals that there is no clear case of *Bd-*GPL consistently possessing more members of a pathogenicity gene family, however there exists intense differences in copy number of pathogenicity genes between *B. dendrobatidis* strains.

We observed phylogenetic differences between the *M36* genes of *B. dendrobatidis* and *P. stewartii*, with the homologs segregated into separate clades by species. Despite the overall similarity in *M36* counts between the two species, the amino acid sequence-level variation was quantified at an average of 49% sequence similarity between *P. stewartii* M36 proteins and their closest homolog in *B. dendrobatidis*. *B. dendrobatidis* contains between 30 and 40 copies of *M36* per strain, while *P. stewartii* has 31. Phylogenetic analysis revealed an ancestral clade of *M36* with copies from *B. dendrobatidis*, *P. stewartii*, *B. salamandrivorans*, and *H. polyrhiza,* along with a clade of *M36* genes exclusive to *B. dendrobatidis*. The abundance of *M36* in *P. stewartii,* together with the ancestral and *B. dendrobatidis*-exclusive gene clades, suggests that *M36* may serve functions unrelated to pathogenicity in most chytrid fungi, while *M36* genes from the *B. dendrobatidis-*exclusive clade may have evolved to enhance *B. dendrobatidis*’s virulence. Although the ancestral *M36* genes were universally present among the *Bd*-BRAZIL and *Bd*-GPL lineages, the *B. dendrobatidis-*exclusive *M36* genes exhibited more variability in presence and absence among the *B. dendrobatidis* genomes. Furthermore, we detected 23 *M36* genes from *Bd*-BRAZIL genomes without homologs in the reference genome JEL423, illustrating the breadth of gene family diversity that is lost while relying upon a single reference genome. Two of the newly discovered *M36* loci in *B. dendrobatidis* were co-localized and shared high sequence similarity with conserved *M36* genes, suggesting that they may have arisen from tandem duplication.

In addition to the overall gains and losses in the putative pathogenicity genes *CBM18*, *M36, ASP, CRN,* and S41 among *B. dendrobatidis* strains, we observe diversity in domain counts within homologs of the CBM18 protein family. The phenomenon of domain gains and losses within a single protein is rare yet has been previously reported in other systems (Prakash & Bateman 2015). In one CBM18 ortholog (BDEG_21734) we note a consistent domain gain from four to five domains between *Bd*-GPL and *Bd*-BRAZIL strains. It is possible that such changes in protein structure could affect the functions of these homologs, however this can not be proven without strenuous functional analysis.

Studies have focused on the genomic repertoire of reference genomes JEL423 and JAM81, however we show that there is a breadth of genomic diversity that is ignored when relying on a single reference genome. Previous work on gene expression variance between *Bd-*BRAZIL and *Bd-*GPL strains has been conducted by aligning transcripts to the JEL423 genome, indicating an up-regulation of some pathogenicity genes in *Bd*-GPL with respect to *Bd*-BRAZIL strains (McDonald et al. 2020). Our transcriptome analysis of CLFT044 transcripts aligned against the CLFT044 and JEL423 genomes suggests that, although mostly accurate, transcription will likely be over or under-estimated for many genes. The results of our differential expression analysis between *Bd*-BRAZIL and *Bd*-GPL lineages suggests that genomic differences (lineage specific gene families and genes with reference dependent transcript counts) could explain ∼12.8% of the DEGs identified. Additionally, we determined that aligning RNA from *Bd*-BRAZIL isolates to the JEL423 assembly will recover ∼2% fewer reads than when aligning to CLFT044. We therefore suggest a candidate reference genome for every *B. dendrobatidis* lineage to more accurately assess future transcriptomic comparisons. While our analysis elucidated a partial pangenome between *Bd*-BRAZIL and the hyper-pathogenic *Bd*-GPL strains, it does not include representatives from *Bd*-CAPE and *Bd*-ASIA lineages. Although there are Illumina genomes for strains from these lineages, high-quality long-read assemblies for these genomes are currently unavailable nor did we have access to those strains for deep sequencing. Our analyses found the Illumina *B. dendrobatidis* assemblies were not comparable to the long-read genomes as they did not capture the full diversity of multi-copy gene families including *M36*, *CRN*, *S41*, *ASP*, or *CBM18*. The absence of homologs from expanded gene families in the short-read genomes, when compared to their long-read counterparts, highlights a significant issue: genome assemblers may collapse multiple, highly similar genes into a singular sequence during assembly of short reads. In contrast, long reads preserve a greater diversity within their flanking sequences, which facilitates a more accurate and comprehensive assembly of the genome’s intricate structure. Additionally, there are currently no long-read genomes for the saprophytic chytrid, *H. polyrhiza,* and efforts to improve the assembly with long-read sequencing may recover missing orthologs from expanded gene families. Future studies including high-quality long-read genomes from all lineages will improve upon the *B. dendrobatidis* pangenome analysis that we report here. We hope these genomic resources will help future exploration of genomic variation in *B. dendrobatidis,* potentially elucidating the mechanisms that make *Bd-*GPL a more globally successful pathogen than the other lineages.

## Materials and Methods

### DNA extraction

Three *B. dendrobatidis* strains from the *Bd-*BRAZIL lineage, CLFT044, CLFT067, and CLFT071, were grown on 1% Tryptone agar for seven days at 21°C. Zoospore and sporangia tissue was harvested from each culture by flooding with 1mL sterile reverse osmosis (RO) water, scraping colonies with an L-spreader, pelleted at 6500g for seven minutes, and flash frozen in liquid Nitrogen. *Ps* JEL0888 was grown in PmTG broth (1 g peptonized milk, 1 g tryptone, 5 g glucose, 1 L distilled water) for 14 days at 23°C before the samples were centrifuged at 6500g for seven minutes to remove broth and flash frozen in liquid Nitrogen. Cetyltrimethylammonium Bromide (CTAB) DNA extraction was performed on the frozen tissue samples from each isolate (Carter-House et al. 2020).

### RNA extraction

Three biological replicates of three *Bd-*BRAZIL and two *Bd*-GPL strains were inoculated onto 1% Tryptone agar plates using 5 x 10^6^ total active zoospores in 2mL of sterile RO water. Cultures were left to incubate under the same conditions as the DNA *B. dendrobatidis* tissue samples until active zoospores were visualized around every colony (5-7 days depending on the isolate). After incubation tissue samples were harvested with an L-spreader and pelleted at 6500g for seven minutes and flash-frozen in liquid nitrogen prior to RNA extraction. Because the samples were not filtered to remove sporangia, tissue samples consisted of both sporangia and zoospores. RNA was extracted from the tissue samples using TRIzol solution (*Invitrogen*, Mulgrave, VIC, Australia) under manufacturer’s protocol coupled with an overnight precipitation in isopropanol at -21°C to increase yields. Total RNA was sent to *Novogene* (Davis, CA) for poly A enrichment, cDNA synthesis and *NovaSeq* PaE150 sequencing to achieve 6Gb data per sample.

### DNA sequencing

gDNA from the three *Bd*-BRAZIL strains was sent to *MiGS* (*SeqCenter*) for Oxford Nanopore Technologies (ONT) sequencing to obtain 900mbp reads (∼30X coverage) for genome assembly. DNA from *Ps* JEL0888 was sequenced with the Oxford Nanopore MinION using the NBD104 barcoding kit and LSK109 ligation sequencing kit following manufacturer’s protocols resulting in 696.915 Mb (∼25X coverage).

### Genome Assembly and Annotation

The ONT reads from all four samples were assembled *de novo* using *Canu* v2.2 (Koren et al. 2017) using the estimated genome size of 25Mb. *Canu* provided scaffolds with telomeres but did not produce the most contiguous assemblies. Assembly was repeated with *MaSuRCA v4.0.9* (Zimin et al. 2013) incorporating publicly available Illumina sequence data for each strain (O’Hanlon et al. 2018; Amses et al. 2022) which produced a more contiguous assembly. The *Canu* assemblies were scaffolded against their respective MaSuRCA assemblies with RAGTAG *v2.1.0* (Alonge et al. 2022) to achieve contiguous genomes with telomeres. The assemblies were polished by 10 iterations of *Pilon* v1.24 (Walker et al. 2014) run within AAFTF (Stajich & Palmer 2022) using publicly available Illumina sequence data (Rosenblum et al. 2013; Farrer et al. 2013; O’Hanlon et al. 2018; Amses et al. 2022; Clemons et al. 2023).

Annotation was performed using Funnanotate v1.8.14 (Palmer & Stajich 2023) utilizing the RNA-seq data from the three *Bd*-BRAZIL strains to increase accuracy of gene predictions for those genomes. Briefly, this entailed sorting the scaffolds by size, RepeatModeller *v2.0.4* (Flynn et al. 2020) and EDTA *v2.1.0* (Ou et al. 2022) to generate a library of transposable elements (TEs) in the *B. dendrobatidis* genome assemblies, RepeatMasker *v4.1.4* to mask the repetitive elements curated by RepeatModeler *v2.0.4* for each assembly, training the *Bd*-BRAZIL assemblies with our RNA-seq data, and functional annotation using default parameters. We downloaded the genome assemblies and annotations for three *Bd-*GPL strains; JEL423 (GCA_000149865), RTP6 (GCA_003595275.1) (Sumpter et al. 2018), and JAM81 (GCF_000203795) (Amses et al. 2022) to compare genomic content between BRAZIL and GPL strains.

We assessed the quality of our genome assemblies using QUAST v5.0.0 (Gurevich et al. 2013) and BUSCO v5.5.0 (Seppey et al. 2019) in genome mode against the fungi_odb10 gene sets. We compared the BUSCO and QUAST results with those from the reference and high quality *Bd-*GPL genomes (supplementary table 1). We calculated telomere counts using pattern searching (with the pattern TAA(C)+) with find_telomeres.py (https://github.com/markhilt/genome_analysis_tools) to assess chromosome completeness for all assemblies. We used PHYling v2.0 (Stajich & Tsai 2023) on the protein annotations for the six *B. dendrobatidis* genomes, *P. stewartii*, *H. polyrhiza* and *B. salamandrivorans* using the fungi_odb10 BUSCO model and default parameters to generate a multi-gene alignment for phylogenetic analysis and confirm relatedness of strains and species. The phylogenetic tree was constructed with RAxML v8.2.12 using the model LG+G8+F and 200 bootstrap replicates (Stamatakis 2014). The final species tree was rendered with ggtree v3.8.2 (Yu 2022).

### Transposable Element Content

We used RepeatModeler *v2.0.4* (Flynn et al. 2020) and EDTA *v2.1.0* (Ou et al. 2022) to generate a library of TEs in the six *B. dendrobatidis* genome assemblies, *P. stewartii, H. polyrhiza,* and the *Batrachochytrium salamandrivorans* (*Bsal*) strain AMP13/1 (GCA_002006685.2) (Wacker et al. 2023). We merged the libraries generated from the different assemblies and collapsed out duplicate TEs using cd-hit *v4.8.1* at 98% sequence identity across the full gene length and default parameters (Huang et al. 2010). We used this condensed TE library and RepeatMasker *v4.1.5* (Smit et al. 2013-2022) to determine counts of LTR, LINE, and DNA TEs in the genomes. Unknown TEs were searched with BLASTN v2.14.1 (Altschul et al. 1997; Camacho et al. 2009) against *RepBase* (Bao et al. 2015) database to classify them as LTR *Copia*, *Gypsy*, or DNA Type II TEs. 882 (38%) of TEs in the condensed library could not be classified to any of the above families.

### Synteny analysis between representative *Bd*-BRAZIL and *Bd*-GPL strains

Scaffold synteny was estimated between the 10 longest scaffolds in *Bd*-BRAZIL strain CLFT044 against the *Bd*-GPL JEL423 *Sanger* and RTP6 *PacBio* genomes using GENESPACE v1.2.0 (Lovell et al. 2022). The syntenic blocks between these three genomes were inferred using default settings, including *Ps* JEL0888 as the outgroup. The GENESPACE v1.2.0 riparian plot was constructed using JEL423 as the reference genome and flipping the orientation of CLFT044 scaffolds; scaffold_1, scaffold_2, scaffold_4, scaffold_5, and scaffold_8 which were inverted with respect to their RTP6 and JEL423 counterparts. We aligned the ONT DNA reads from CLFT044 back against the CLFT044 genome using minimap2 v2.24 (Li 2018) to confirm that reads supported merged scaffolds observed in CLFT044 but not JEL423 or RTP6.

### Gene Family Variation between B. dendrobatidis and related species

We used Orthofinder v2.5.4 (Emms & Kelly 2019) as an initial step towards comparative genomics by identifying gene families unique to the *Batrachochytrium* genus (*B. dendrobatidis* and *B. salamandrivorans*), but absent in their saprophytic relatives. This included the three *Bd-*BRAZIL ONT-hybrid genomes and *P. stewartii,* the three public *Bd*-GPL genomes; JEL423, JAM81, and the published PacBio *B. salamandrivorans* genome APM13/1, and the Illumina genome for *H. polyrhiza* Gene IDs of JEL423, belonging to gene families specific to the pathogenic chytrids *B. dendrobatidis* and *B. salamandrivorans*, were searched for Gene Ontology (GO) terms against the FungiDB database, which assigns GO terms based on protein domains and manual user contributions that are regularly reviewed and updated. Additionally we used Orthofinder to identify gene content differences between the three *Bd*-GPL and three *Bd-*BRAZIL long-read genomes to identify the core and pangenome of *B. dendrobatidis*. Orthofinder visualizations were rendered with UpsetR v1.4.0 (Conway et al. 2017). We incorporated the species tree generated with phyling and the Orthofinder results to assess gene family expansions and contractions with CAFE v5.0.0 (Mendes et al. 2021). CAFE visualizations were rendered with CafePlotter v.0.2.0 https://github.com/moshi4/CafePlotter.

### Presence Absence Variation (PAV) of Pathogenicity genes in *B. dendrobatidis* and other chytrids

We conducted HMMsearch 3.3.2 with an e-value threshold of 1e-15 for PFAMs PF02128, PF03572, PF00026, and PF00187 which included 7K M36, 100K S41, 58K ASP, and 11K CBM18 proteins, respectively, to quantify the number of these proteins in each annotated genome (Eddy, 2011). HMMsearches using the Crinkler necrosis (CRN) PFAM PF20147 were unsuccessful at detecting copies in *B. dendrobatidis*. We used HMMbuild on the previously 106 identified CRN proteins in JEL423 to construct an hmm profile to screen and count CRN proteins in *B. dendrobatidis* strains (Farrer et al. 2017). We constructed a heatmap for the protein family counts using the R package pheatmap v.1.0.12 (Kolde 2019). Alignments were performed using MUSCLE *v5.1* (Edgar 2004) prior to tree building and downstream analysis.

Since orthologous genes are likely to be syntenic with orthologs from closely related strains, we used cblaster *v1.3.18* (Gilchrist et al. 2021) with a minimum percent identity of 80% and two flanking genes on the 5’ and 3’ end to identify the syntenic homologs of every *M36* and *CBM18* across the *B. dendrobatidis* strains. Genes were required to share the same four flanking genes, two upstream and two downstream of the *M36* or *CBM18*, to be considered orthologous. This analysis identified the non-redundant set of orthologous *CBM18* or *M36* across all strains, preferentially using the JEL423 copies given its status as a primary reference genome. If orthologs were missing in JEL423, we represented them by a copy from another strain. We used MUSCLE *v5.1* (Edgar 2004) to align the non-homologous *M36* and *CBM18* encoding genes and constructed phylogenetic trees for both gene sets with IQTREE2 *v2.2.2.6* (Minh et al. 2020) using the ModelFinder function which selected the VT+R10 model for *CBM18* and the TWM+F+R5 model for *M36* with 1000 SH-like bootstrap replicates (Kalyaanamoorthy et al. 2017). We included the *M36* and *CBM18* genes from *H. polyrhiza* and *P. stewartii* as outgroups and to root the phylogenies. Phylogenetic visualizations were rendered with the R package, ggtree v3.8.2 (Yu 2022).

CBM18 domain-containing proteins have been reported to contain multiple domains per gene in *B. dendrobatidis* strain JEL423 (Abramyan & Stajich 2012; Farrer et al. 2017). The diversity of JEL423’s CBM18 repertoire includes variable counts of Tyrosinase and Deacetylase domains, as well as Lectin-like CBM18 proteins without additional domains (Abramyan & Stajich 2012). We used HMMSCAN to catalog the variation in CBM18 domain copy number between *CBM18* homologs in the *B. dendrobatidis* Long-read genomes (Eddy 2011).

### RNA-seq Read Mapping

Based on the genomic diversity we observed between *B. dendrobatidis* strains, we questioned the efficacy of using a single reference genome for transcriptomic analysis of different *B. dendrobatidis* strains. To assess whether a *Bd*-BRAZIL reference genome will increase recovery of *Bd*-BRAZIL transcripts, we aligned RNA-seq reads from four *Bd-*BRAZIL and two *Bd*-GPL strains were aligned to the *Bd*-BRAZIL CLFT044 assembly and to the *Bd*-GPL genome for JEL423 (GCA_000149865) using HISAT2 *v2.2.1* (Kim et al. 2019). All sequence reads were mapped to the indexed genomes. Raw read counts were generated with FeatureCounts from Rsubread (Liao et al. 2019) and TPM values were calculated with edgeR (Robinson et al. 2010). Given the genetic variation between *B. dendrobatidis* strains, we tested how much the reference genome matters when calculating gene expression in *B. dendrobatidis* RNA-seq data. We aligned the three replicates of paired end RNA-seq reads from CLFT044 against the CLFT044 genome and against the JEL423 genome to calculate the ratio of TPMs when aligning to CLFT044 (self) over the reference genome JEL423 (GCA_000149865). We focused this analysis on Single-Copy orthologous gene families, genes determined from Orthofinder to be single copy in both genomes, to avoid ambiguity caused by comparing distant orthologs. Using a global-pairwise sequence alignment of the two orthologous coding sequences (Pearson 2000) we scored sequence pairs for their alignability and then evaluated the relationship of sequence divergence, number of secondary blast hits, gap openings, and mismatch differences between CLFT044 and JEL423 to test whether length or sequence differences between homologous genes explain the variation in TPM calculations.

We tested whether genomic differences have inflated the number of differentially expressed genes (DEGs) between *Bd-*BRAZIL and *Bd-*GPL strains. A previous study reported DEGs between the *Bd-*BRAZIL and *Bd-*GPL lineages when RNA from two *Bd-*BRAZIL strains (CLFT044 and CLFT001) and four *Bd-*GPL strains (CLFT023, CLFT026, JEL410, and JEL422) are aligned to the reference genome strain JEL423 (McDonald et al. 2020). We performed HISAT2 alignments of the RNA-seq data from this study to the reference genome of strain JEL423 and separately aligned to CLFT044. We used the R packages FeatureCounts and DeSeq2 *v*1.4.2 (Love et al. 2014) to re-calculate the number of differentially expressed genes (log2fold ≥ 1.5 and Bonferroni adjusted p-value ≥ 0.05) between the *Bd-*BRAZIL and *Bd-*GPL lineages. We intersected the identities of differentially expressed genes with the identities of *Bd-*GPL /*Bd*-BRAZIL specific genes from Orthofinder and the identities of the single copy orthologous genes with reference dependent transcript counts. Images depicting this intersection were rendered using the R package ggvenn *v.* 0.1.10 (Yan 2023).

## Acknowledgements

JES is a Fellow in CIFAR program Fungal Kingdom: Threats and Opportunities. The work was partially supported by a catalyst grant from CIFAR and CIFAR fellowship funds and U.S. Department of Agriculture, National Institute of Food and Agriculture Hatch projects CA-R-PPA-211-5062-H. The Gordon and Betty Moore Foundation Award #9337 (<u>10.37807/GBMF9337</u>) to Lilian Fritz-Laylin (PI), Timothy Y James, and Jason Stajich supported Mark Yacoub. Genome assembly and annotation were performed on the IIGB High-Performance Computing Cluster supported by NSF DBI-1429826, DBI-2215705, and NIH S10-OD016290 grants. We thank Dr. Timothy Y. James and the culture contributors of the Collection of Zoosporic EuFungi at University of Michigan (CZEUM) for providing the *Bd*-BRAZIL and *P. stewartii* isolates used in this study. We would also like to thank Dr. Timothy Y. James, Dr. Cassie Ettinger, Dr. Tania Kurbessoian, Dr. Jessica Huang, Kian Kelly, and Julia Adams for helpful suggestions on this manuscript.

## Author Contributions

M.N.Y. and J.E.S. designed research; M.N.Y. and J.E.S. performed research; M.N.Y. and J.E.S. contributed new data/reagents/analytic tools; M.N.Y. analyzed data; J.E.S. provided supervision and funding; and M.N.Y. wrote the paper with editing by J.E.S.

## Data Availability

The primary sequence data for Nanopore and Illumina DNA sequencing data are under BioProjects PRJNA987700 (*Polyrhizophydium stewartii* JEL0888), and *B. dendrobatidis* strains PRJNA987741 (CLFT067), PRJNA821523 (CLFT044), and PRJNA913953 CLFT071. RNA sequencing data are deposited under the accession numbers GSE253912 and GSE246809. Genome assemblies for *B. dendrobatidis* strains, CLFT044, CLFT067, and CLFT071 are deposited under the Accession numbers; GCA_036783925.1, GCA_036289345.1, and GCA_029704095.1 respectively. An updated genome assembly of *P. stewartii* was generated as part of this publication and deposited as GCA_027604665.2. The RTP6 genome assembly previously published was updated with gene annotation from this study and deposited as accession GCA_003595275.2. The TE sequences and annotations can be downloaded from the github repository for this project at **DOI 10.5281/zenodo.15070156**

