## Supplemental Figures and Tables for "Comparative genomics reveals intra and inter species variation in the pathogenic fungus *Batrachochytrium dendrobatidis*"

| | CLFT044 | CLFT067 | CLFT071 | JAM81 | JEL423 | RTP6 | JEL0888 | JEL142 |
|---|---|---|---|---|---|---|---|---|
| *Scaffold Count* | 79 | 86 | 85 | 127 | 70 | 63 | 291 | 11986 |
| *BUSCO Assembly* | 88.9 | 88.8 | 88.9 | 88.9 | 88.2 | 89.2 | 81.4 | 80.9 |
| *BUSCO Annotation* | 92 | 91.2 | 91.7 | 88.8 | 81.5 | 92.1 | 87.9 | 62.7 |
| *Total Length* | 26117516 | 22029792 | 22218474 | 24315081 | 23897668 | 24102166 | 31203426 | 26111666 |
| *N50* | 1102615 | 1138468 | 953307 | 1484462 | 1707251 | 1511395 | 350379 | 45230 |
| *N90* | 136200 | 282475 | 258354 | 561733 | 857155 | 361838 | 77712 | 1123 |
| *L50* | 7 | 7 | 9 | 6 | 5 | 6 | 25 | 138 |
| *L90* | 28 | 19 | 25 | 17 | 14 | 17 | 97 | 1123 |
| *Ns per 100kbp* | 6.7 | 14.55 | 10.22 | 759.89 | 1338.77 | 0 | 0 | 933.95 |
| *Largest Contig* | 4271105 | 2486897 | 1945879 | 4429270 | 4440149 | 4481161 | 1456518 | 227101 |
| *Telomeres TOTAL* | 26 | 0 | 6 | 0 | 0 | 7 | 1 | 3 |
| *Telomeres Forward* | 15 | 0 | 3 | 0 | 0 | 6 | 0 | 2 |
| *Telomeres Reverse* | 11 | 0 | 3 | 0 | 0 | 1 | 1 | 2 |
| *Telomere to Telomere Scaffolds* | 3 | 0 | 0 | 0 | 0 | 0 | 0 | 0 |

**Supplementary Table S1:** Genome assembly statistics for the genomes used in this study.

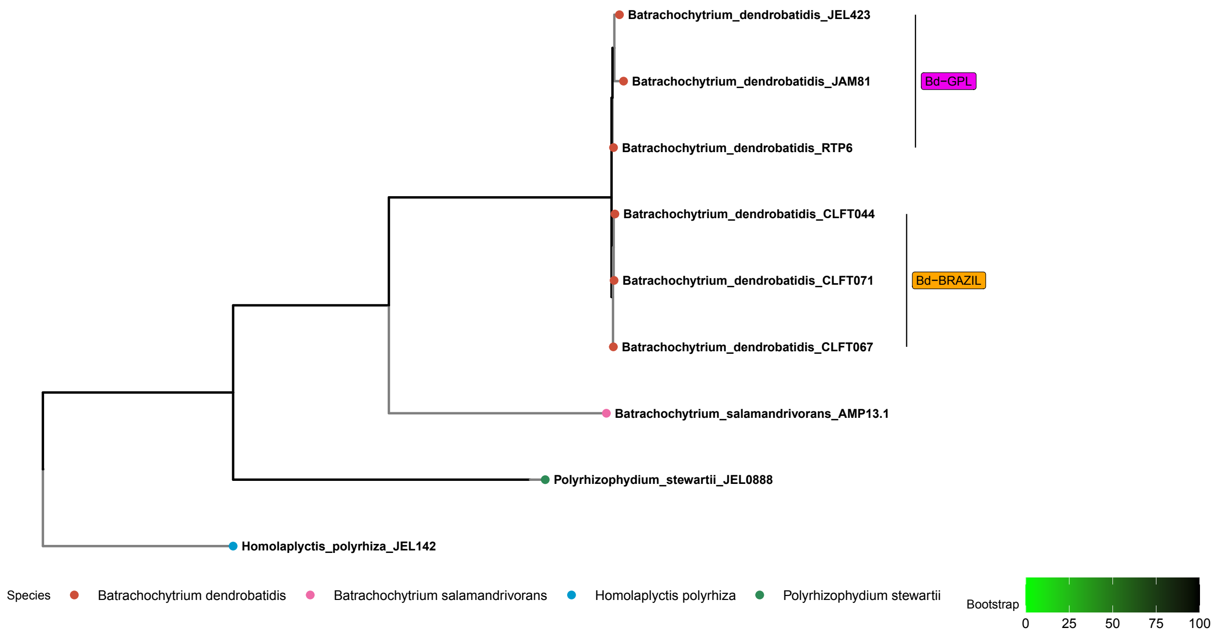


**Supplementary figure S1.** Multi-gene phylogeny inferred by phyling of genomes used in this study. Tree tips are color-coded by species. *Bd*-GPL and *Bd*-BRAZIL lineages are indicated by clade labels.

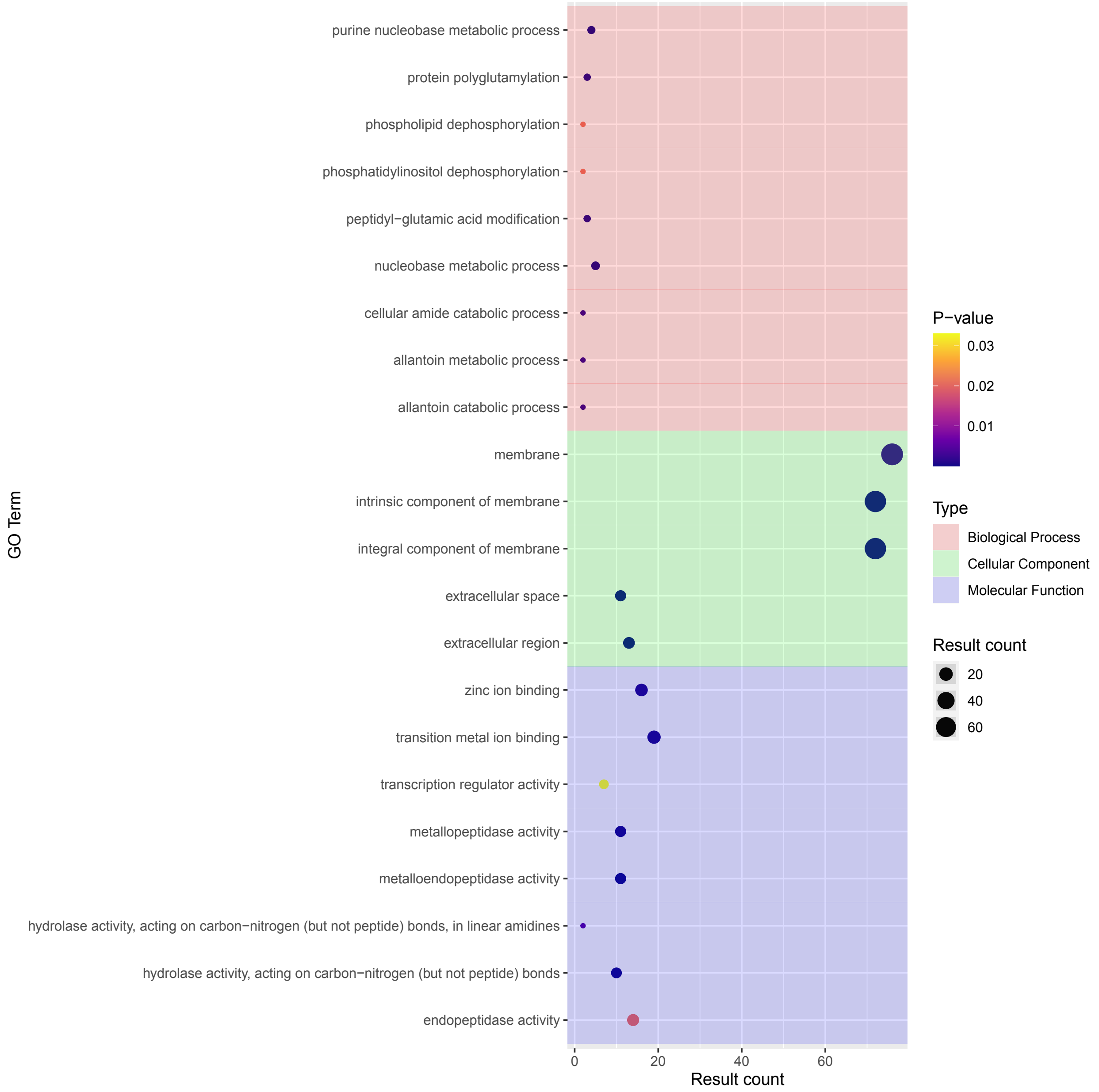


**Supplementary figure S2. Results of Gene Ontology (GO) enrichment for the gene families specific to *Bsal* and *Bd* but absent in *Ps* and *Hp*.** GO terms for the pathogen specific gene families are shown for the three classes; Biological process, cellular component, and molecular function. The Y-axis lists the REVIGO filtered GO terms. The X-axis and bubble size reflects the number of genes within the pathogen specific orthogroups from *Bd* strain JEL423 associated with that GO term.

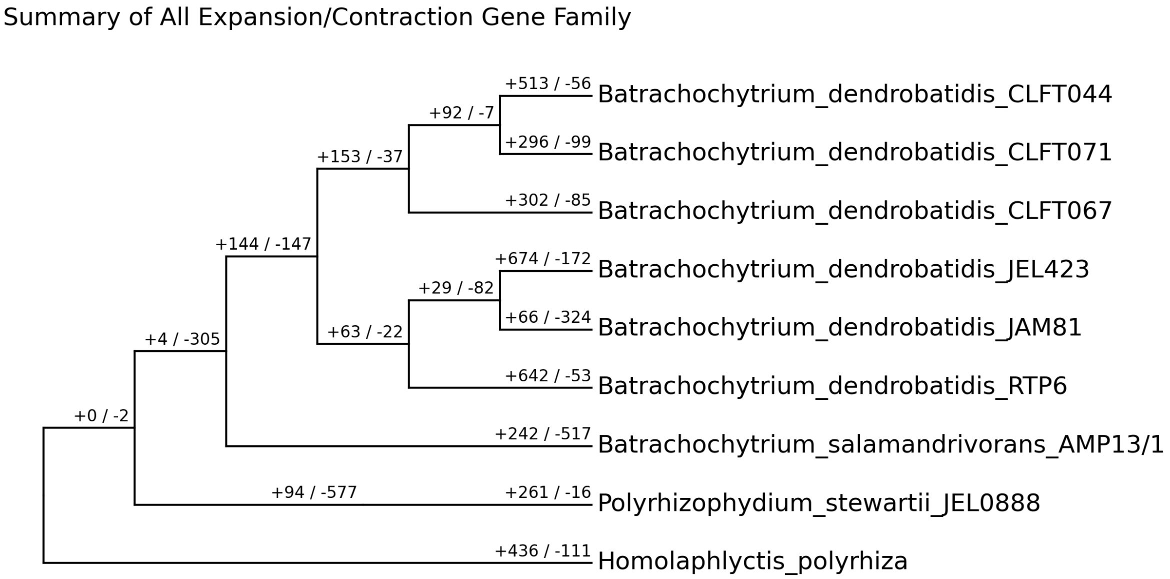


**Supplementary Figure S3: Lineage Specific gene gains and losses in *Hp*, *Ps*, *Bd*, and *Bsal*.** An ultrametric phylogeny of *Bd* and related chytrids generated with Phyling v2.0 is shown. Values on the branches indicate gene family expansions and contractions. Expansions are indicated with positive values and contractions are represented by negative values.

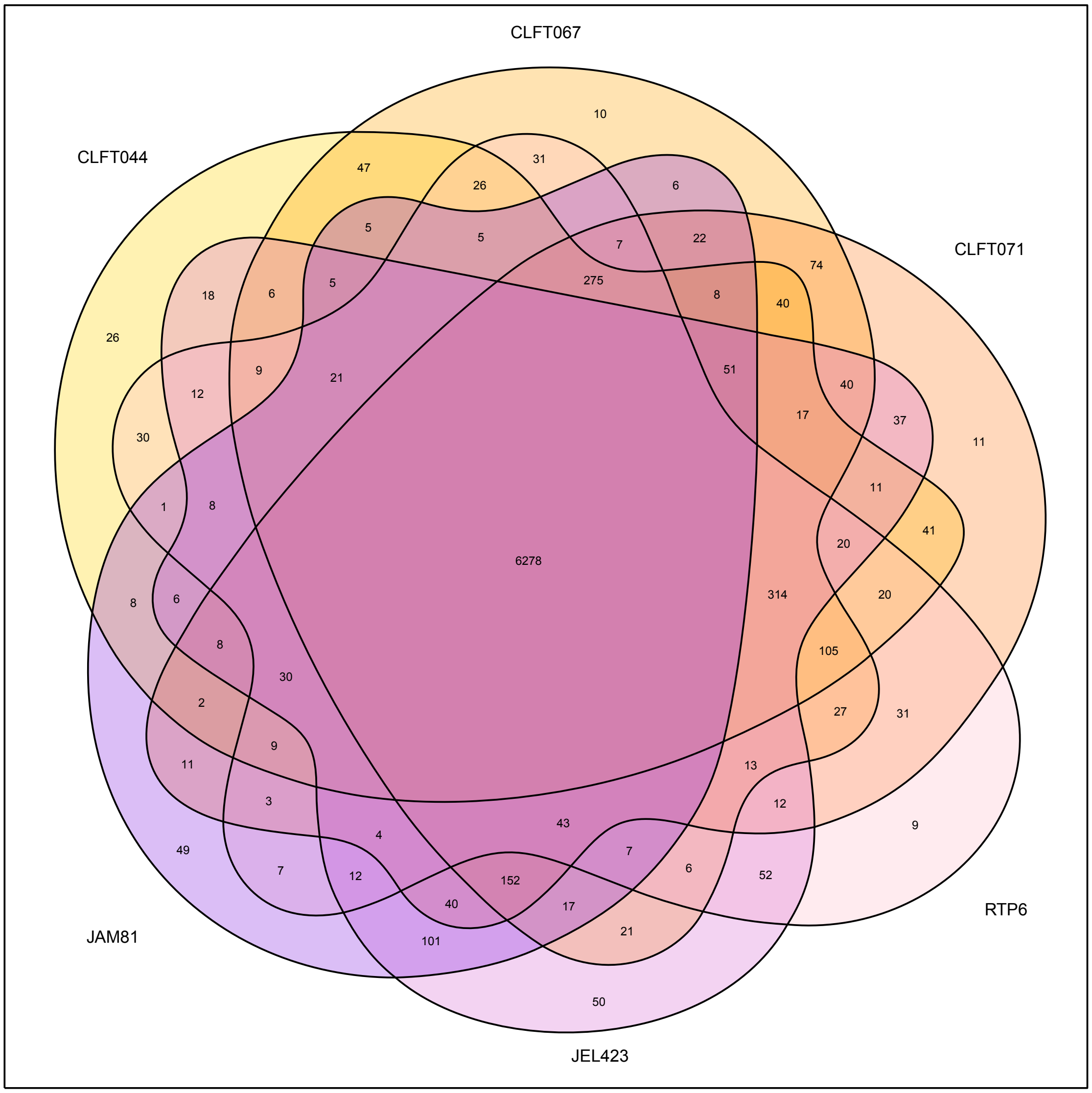


**Supplementary figure S4. Venn diagram depicting the number of core, dispensary, and singleton gene families in the *Bd* pangenome.** Orthology was determined from the Orthofinder results. Numbers represent the counts of gene families within each group.

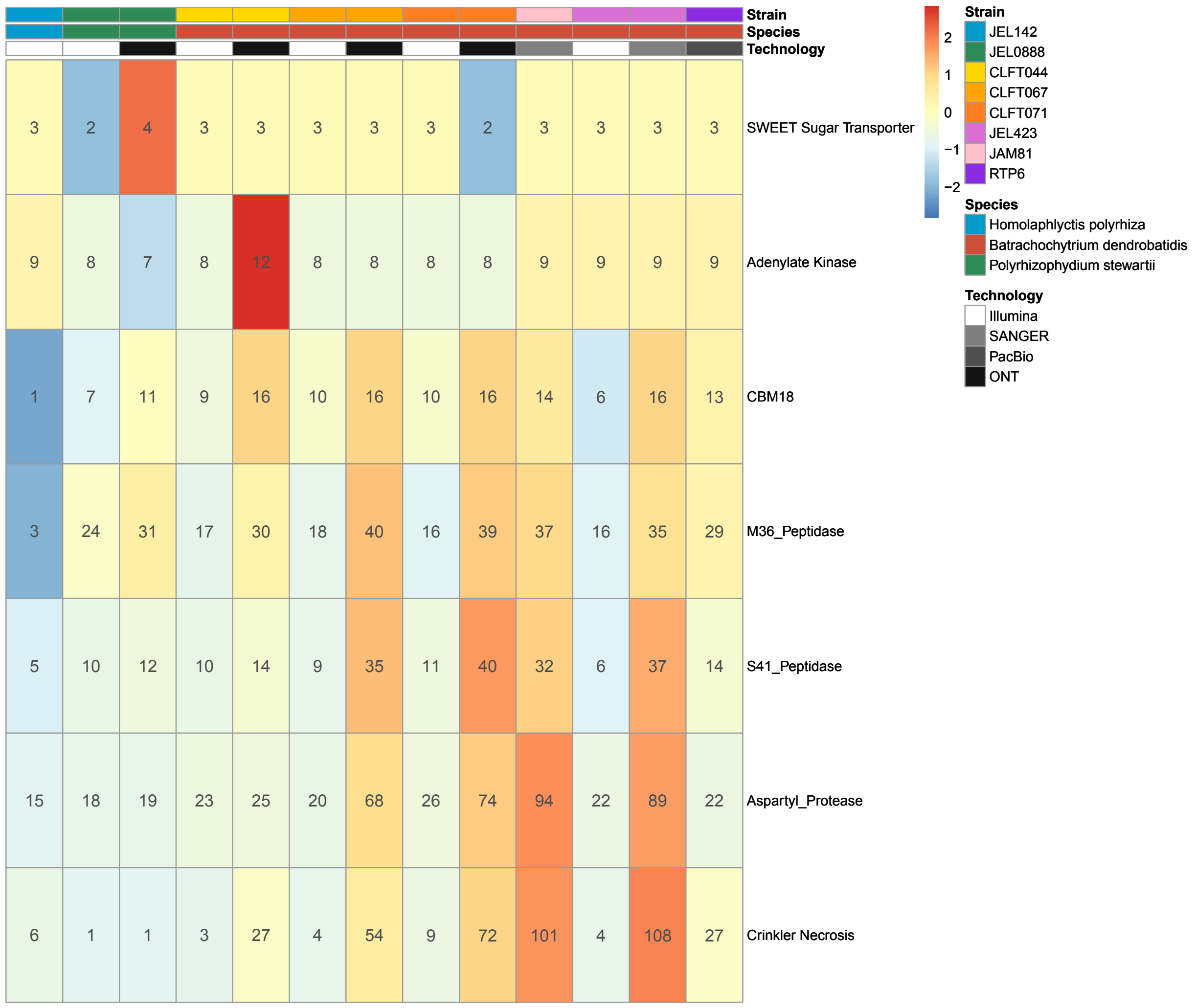


**Supplementary figure S5: Copy number variation for pathogenicity genes *CBM18, M36, S41, ASP*, and *CRN* between *Bd* and saprophytic chytrids.**

Heatmap depicting the copy number variation of pathogenicity genes (SWEET and Adenylate Kinase included as controls). The colors of the gene counts are normalized per gene family with red and blue indicating high and low counts respectively. The sequencing technology used to produce each genome is shown by the gray-scale bar while the top color bar indicates the species for each strain.

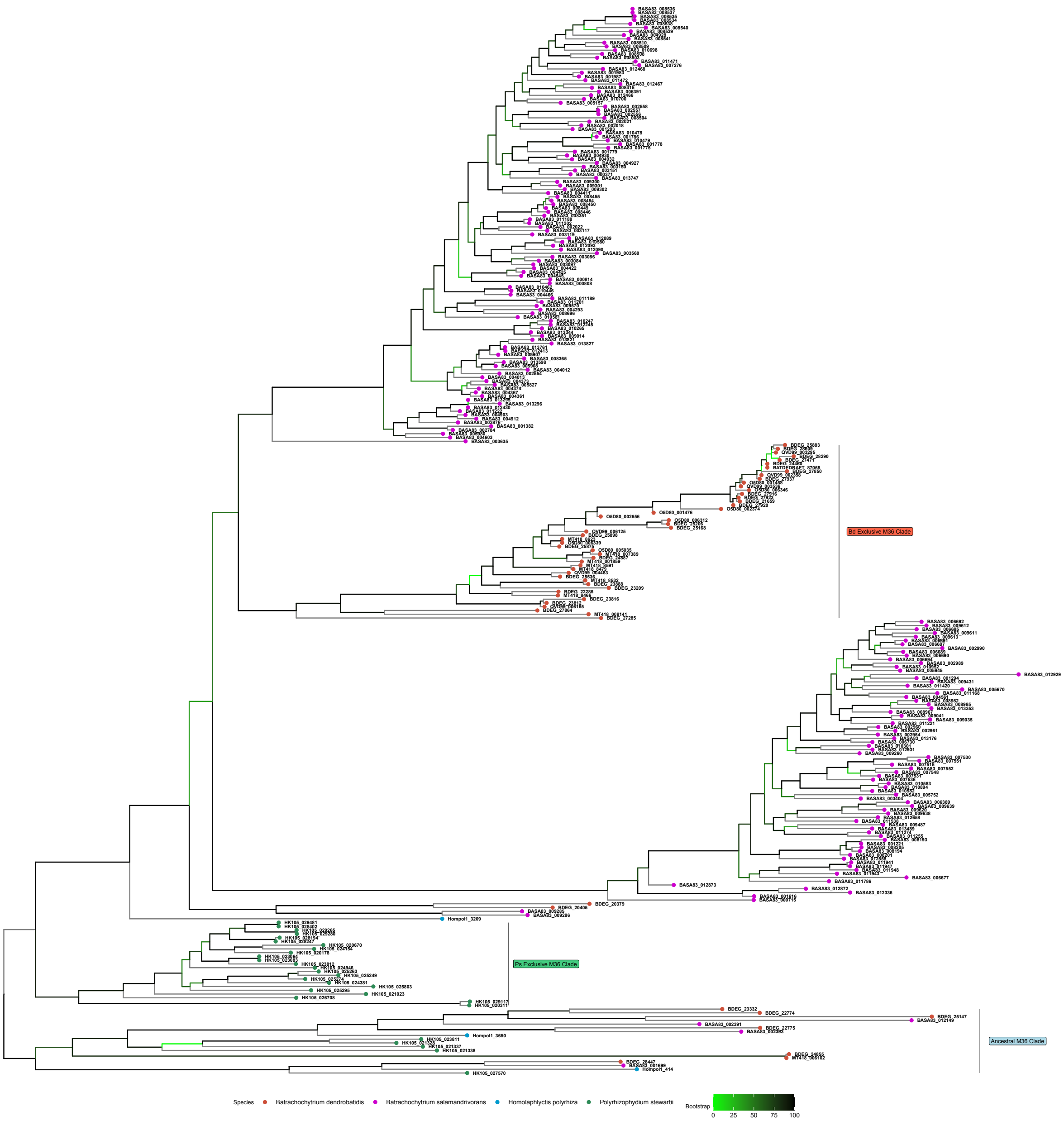


**Supplementary figure S6. Phylogeny of *M36 Peptidase* coding sequences from *Bd*, *Bsal*, *Hp*, and *Ps*.** The *Bd* specific, *Ps* specific, and Ancestral M36 clades are labeled. Tree tips are color-coded by species. Branches are color-coded by bootstrap values.

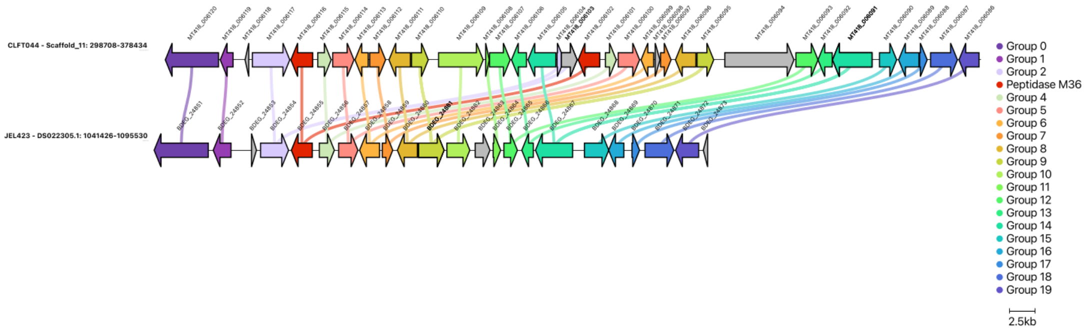


| | BDEG_24851 | BDEG_24852 | BDEG_24853 | BDEG_24854 | BDEG_24855 | BDEG_24856 | BDEG_24857 | BDEG_24858 | BDEG_24859 | BDEG_24860 | BDEG_24861 | BDEG_24862 |
|---|---|---|---|---|---|---|---|---|---|---|---|---|
| Ortholog | MT419_006120 | MT419_006119 | NONE | MT419_006117 | MT419_006116 | MT419_006115 | MT419_006114 | MT419_006113 | MT419_006112 | MT419_006111 | MT419_006110 | MT419_006109 |
| Paralog | NONE | NONE | NONE | MT418_006104 | MT418_006102 | MT418_006101 | MT418_006100 | MT418_006099 | MT418_006097 | MT418_006096 | MT418_006095 | NONE |

| | BDEG_24863 | BDEG_24864 | BDEG_24865 | BDEG_24866 | BDEG_24867 | BDEG_24868 | BDEG_24869 | BDEG_24870 | BDEG_24871 | BDEG_24872 | BDEG_24873 |
|---|---|---|---|---|---|---|---|---|---|---|---|
| Ortholog | NONE | MT419_006108 | MT419_006107 | MT419_006106 | MT419_006105 | NONE | MT419_006090 | MT419_006089 | MT419_006088 | MT419_006087 | MT419_006086 |
| Paralog | NONE | NONE | NONE | NONE | NONE | NONE | NONE | NONE | NONE | NONE | NONE |

**Supplementary figure S7. Evidence of tandem duplications as a mechanism for *M36* gene expansion.** A clinker synteny comparison between CLFT044 and JEL423 depicting tandem duplication in this region. Linkages indicate orthologous genes. Red arrows indicate the *M36* loci. The included table names the JEL423 gene (BDEG locus tags) and their CLFT044 ortholog or paralog.

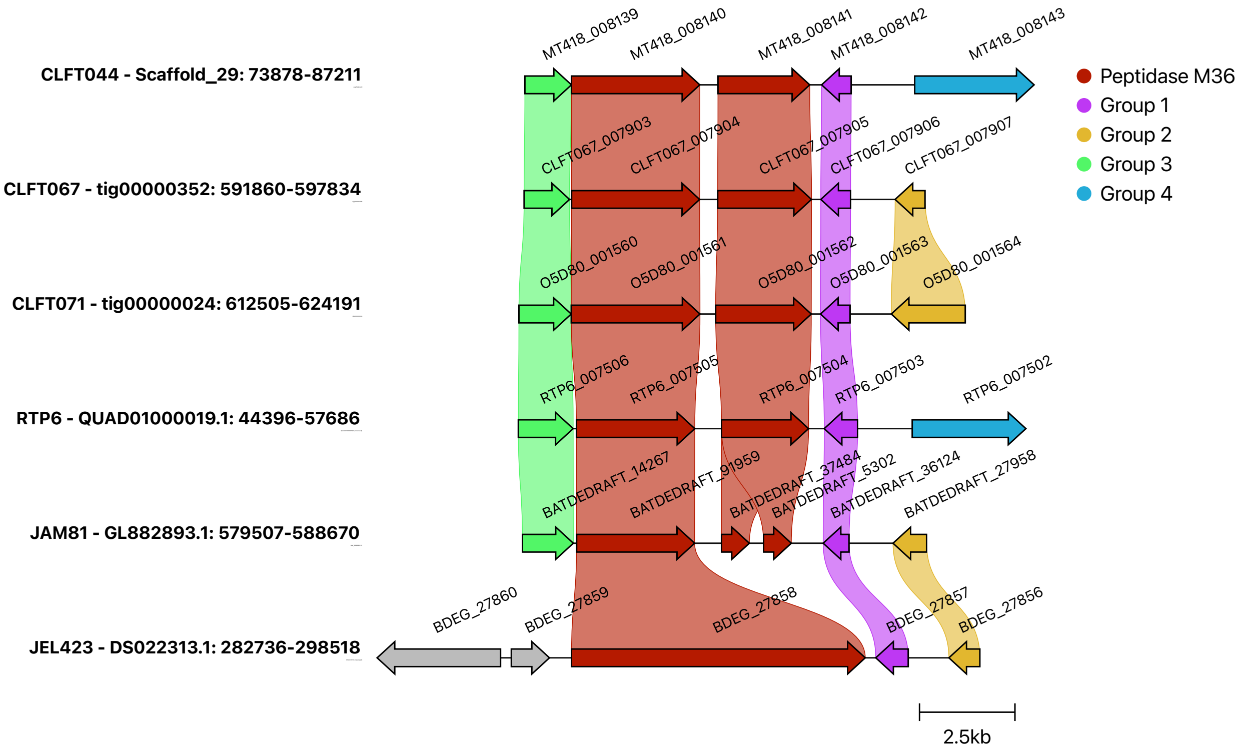


**Supplementary figure S8. Evidence of annotation errors in GPL reference genomes reveal new *M36* loci.** Clinker synteny comparison between regions containing *M36* gene, BDEG_27858 in JEL423 against the other long-read genomes. Linkages indicate regions of  sequence similarity. The region contains two tandemly duplicated *M36* genes in the *Bd*-BRAZIL and RTP6 strains while these have been merged into a single locus in JEL423 and broken in JAM81.

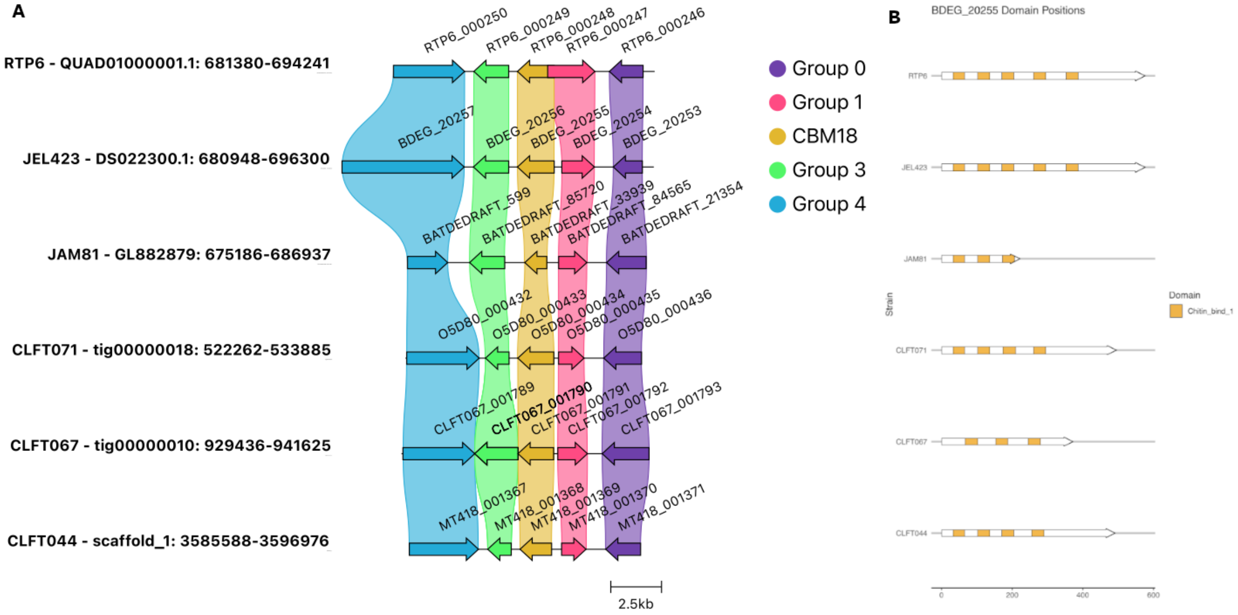


**Supplementary figure S9. Structural differences in the CBM18 homologs of BDEG_20255 (yellow) between different strains of *Bd*.** A) Synteny comparison between the orthologs of BDEG_20255 across *Bd* strains and their neighboring genes, indicating homology of these proteins. B) Structural variation between the BDEG_20255 orthologs in the *Bd* long-read genomes. Different strains display variable counts of CBM18 domains in the same protein.

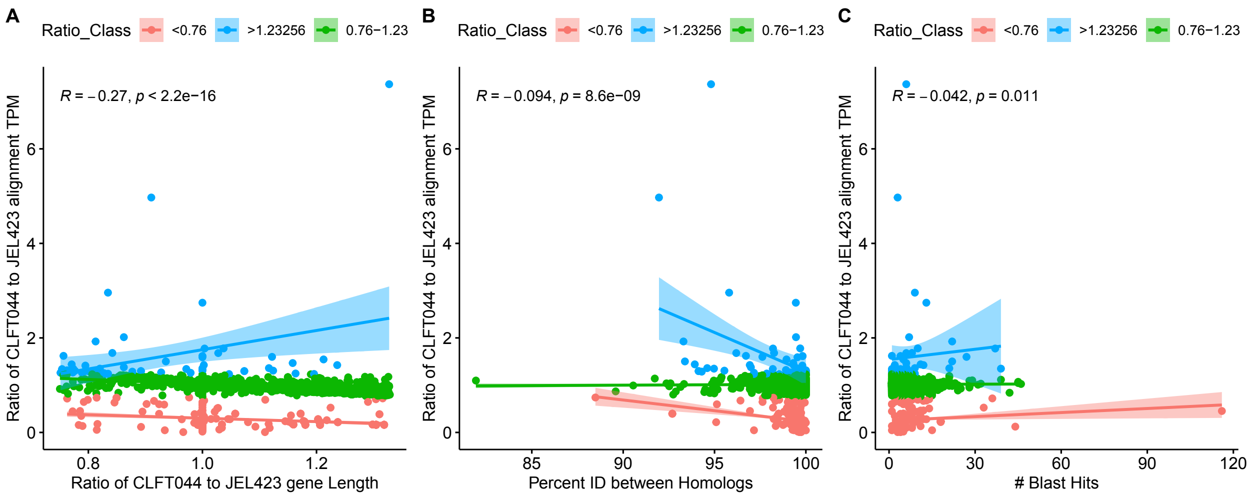


**Supplementary figure S10. Length differences, number of secondary hits, and sequence differences between homologous genes in the self genome (CLFT044) and the reference genome (JEL423) are correlated with TPM ratio**. We tested whether differences in (A) gene length (B) percent identity (C) and number of secondary tblastn hits in the genomes explained why transcripts from many single copy orthologs are mis-counted (TPM ratios above and below 1) when aligning CLFT044 transcripts against the JEL423 genome rather than the CLFT044 genome.

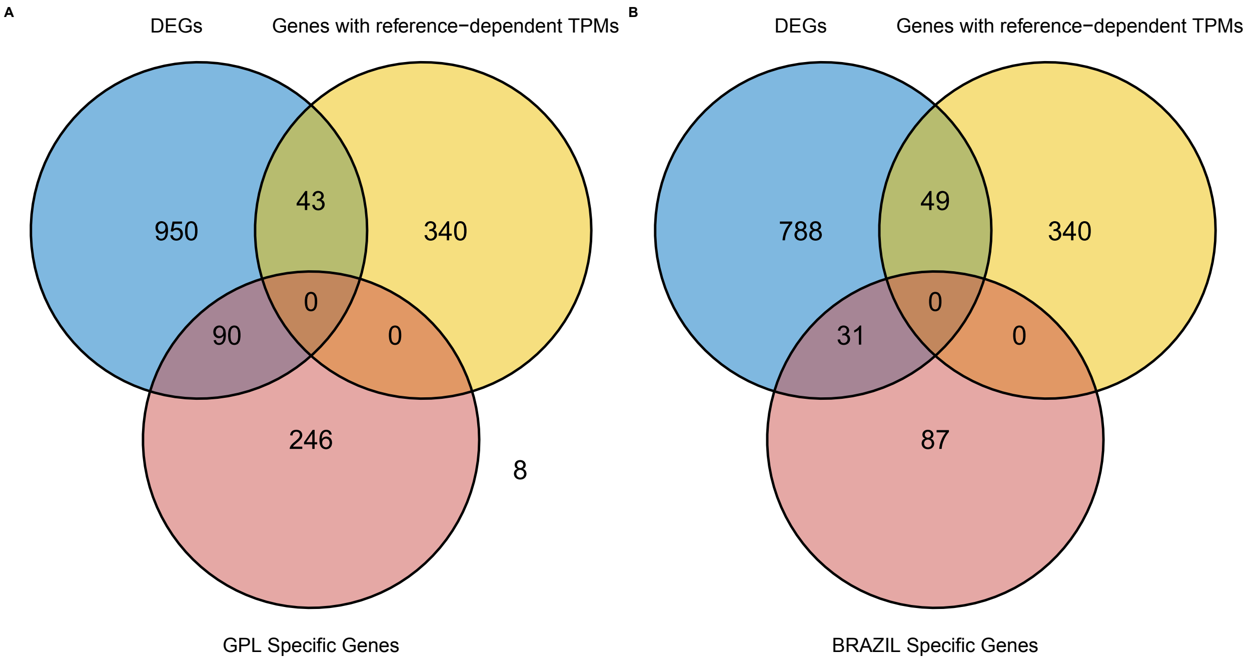


**Supplementary figure S11. The intersection of differentially expressed genes between *Bd* lineages with lineage specific genes and genes with reference dependent transcript counts.** (A) The intersection of differentially expressed genes between *Bd*-GPL and *Bd*-BRAZIL using JEL423 as the reference alignment genome (blue) with the *Bd*-GPL specific genes (red), and genes with reference dependent transcript counts (yellow). (B) ) The intersection of differentially expressed genes between *Bd*-GPL and *Bd*-BRAZIL using CLFT044 as the reference alignment genome (blue) with the *Bd*-BRAZIL specific genes (red), and genes with reference dependent transcript counts (yellow).
